# Nanoscale structural organization and stoichiometry of the budding yeast kinetochore

**DOI:** 10.1101/2021.12.01.469648

**Authors:** Konstanty Cieslinski, Yu-Le Wu, Lisa Nechyporenko, Sarah Janice Hörner, Duccio Conti, Michal Skruzny, Jonas Ries

## Abstract

Proper chromosome segregation is crucial for cell division. In eukaryotes, this is achieved by the kinetochore, an evolutionarily conserved multi-protein complex that physically links the DNA to spindle microtubules and takes an active role in monitoring and correcting erroneous spindle-chromosome attachments. Our mechanistic understanding of these functions and how they ensure an error-free outcome of mitosis is still limited, partly because we lack a comprehensive understanding of the kinetochore structure in the cell. In this study, we use single-molecule localization microscopy to visualize individual kinetochore complexes *in situ* in budding yeast. For major kinetochore proteins, we measured their abundance and position within the metaphase kinetochore. Based on this comprehensive dataset, we propose a quantitative model of the budding yeast kinetochore. While confirming many aspects of previous reports based on bulk imaging, our results present a unifying nanoscale model of the kinetochore in budding yeast.

## Introduction

Cell division is a process of paramount importance for organismal life, ultimately ensuring the faithful propagation of the genome in space and time. Erroneous chromosome segregation can lead to aneuploidy, where daughter cells receive an aberrant karyotype which, in turn, may result in developmental defects or cell death (Santaguida and Amon, 2015). A multiprotein complex called kinetochore assembles at the centromere of each sister chromatid to generate robust connections between chromosomes and spindle microtubules (reviewed in (Musacchio and Desai, 2017)). The general architecture of the kinetochore is conserved in all eukaryotes (Drinnenberg et al., 2016; Hooff et al., 2017). A simple model to study its properties is the budding yeast, *Saccharomyces cerevisiae*, where the kinetochore assembles onto one nucleosome and is attached to one microtubule (Winey et al., 1995). Conversely, multiple copies of units analogous to the budding yeast kinetochore bind to many microtubules in other fungi and multicellular organisms (Zinkowski et al., 1991; Musacchio and Desai, 2017). The kinetochore takes part in several processes during mitosis including maintaining proper chromosome attachment to the spindle, translating the pushing-pulling forces into chromosome movement and controlling the mitotic progression through the spindle assembly checkpoint (Aravamudhan et al., 2015; Asbury, 2017; Joglekar et al., 2010). These functions are strongly dependent on the kinetochore’s structure and its potential remodeling over the cell cycle (Conti et al., 2017; Dhatchinamoorthy et al., 2017; Joglekar et al., 2009).

Early electron microscopy studies defined three electron-dense regions in the kinetochore—the inner kinetochore, the outer kinetochore, and the fibrous corona (Rieder, 1982). In *S. cerevisiae*, where the corona is absent, the inner kinetochore includes the centromeric nucleosome containing an H3 variant called Cse4, the CBF3 complex (Cep3, Ndc10, Ctf13, Skp1), the Mif2 and Cnn1 module (Cnn1, Ctf3, Wip1, Mcm16/22, Mhf1/2), Nkp1/2, the COMA complex (Ctf19, Okp1, Mcm21, Ame1), and Chl4/Iml3. The outer kinetochore consists of the microtubule-interacting network built by Spc105, the MIND complex (Mtw1, Dsn1, Nnf1, Nsl1), the Ndc80 complex (Ndc80c; Ndc80, Spc24, Spc25, Nuf2) and the Dam1 complex (Dam1c) ring (Musacchio and Desai, 2017; Figure 1A with human counterparts shown in the upper right corner of each protein).

**Figure 1:**
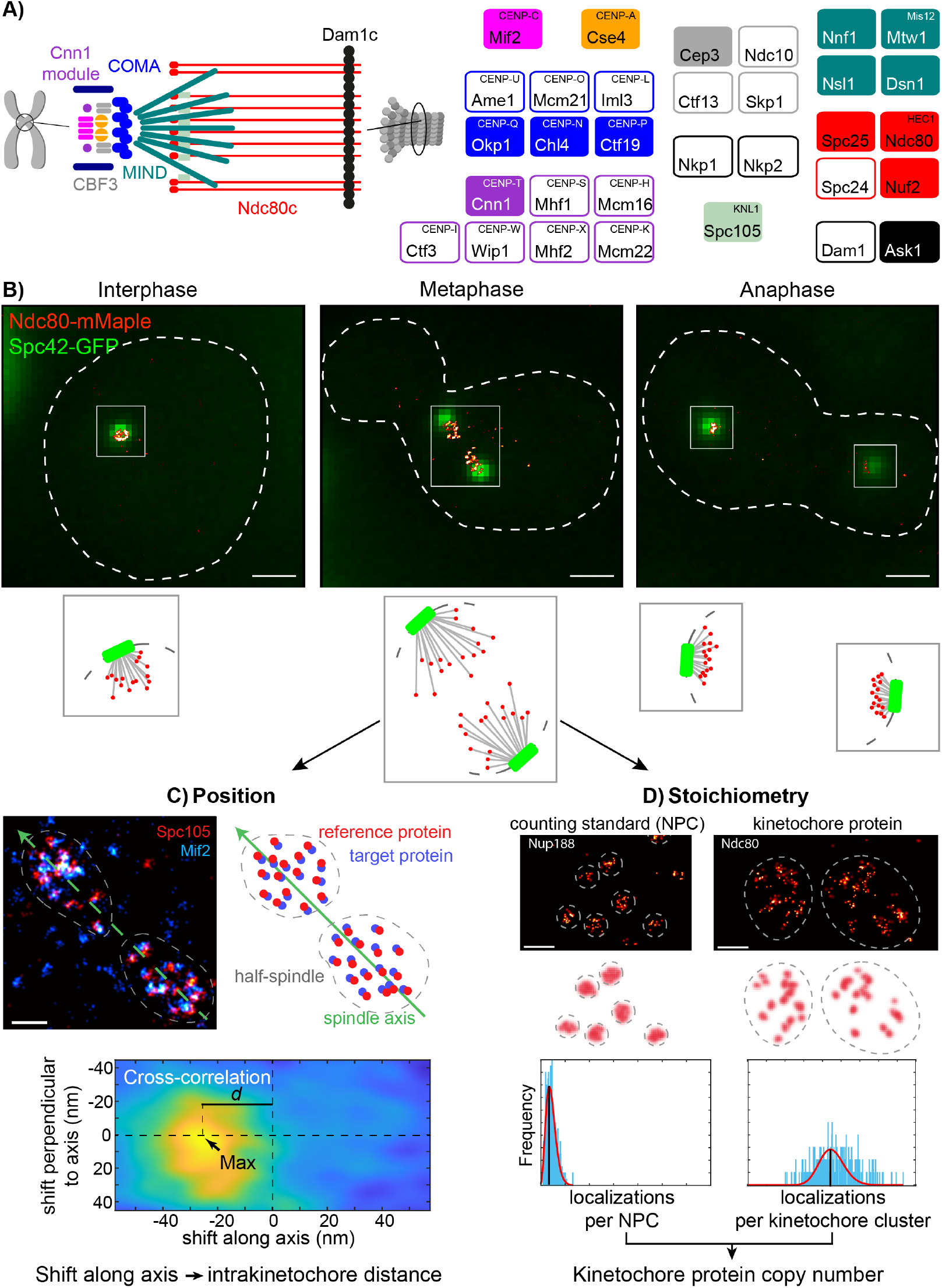
Overview of the study. **A. Protein composition of the budding yeast kinetochore.** Kinetochore proteins are grouped and color-coded by subcomplexes. Only opaquely colored components were measured in this study. Human counterparts are shown in a superscript. **B. Example kinetochore clusters**. Overlays of representative super-resolved images of the kinetochore protein Ndc80 (red) and the diffraction-limited spindle pole body protein Spc42 (green) at different stages of the cell cycle and corresponding cartoons of the budding yeast spindles. Scale bars: 1 μm. **C. The position of kinetochore proteins along the spindle axis**. We labeled and imaged always the reference protein Spc105 (red) together with the target protein (cyan, Mif2 in this example). We manually segmented single kinetochore clusters, defined the spindle axis and calculated the image cross-correlation. The position of the cross-correlation peak corresponds to the average distance between reference and target proteins in the half spindle. **D. Stoichiometry of the budding yeast kinetochore**. We quantified the copy numbers of kinetochore proteins using the nuclear pore complex (NPC) component Nup188, which has 16 copies per NPC, as a counting reference standard. In each experiment, we mixed two strains in which either Nup188 or the target kinetochore protein were labeled with the same fluorescence protein tag mMaple. We then imaged both strains simultaneously. We calculated the ratio of mean localization counts per structural unit (either NPC or kinetochore cluster) between the two proteins. From the relative number of localizations and the known stoichiometry of Nup188, we computed the copy number of the target kinetochore protein. Scale bars: 200 nm.

Despite advances in the last decades in understanding kinetochore composition, a complete picture of its organization in cells is still unclear. A significant portion of the components of both human and budding yeast kinetochores have been already crystallized or analyzed by cryo-EM (for an overview see Dimitrova et al., 2016; Hinshaw and Harrison, 2019; Jenni and Harrison, 2018; Musacchio and Desai, 2017; Yan et al., 2019). Most of the structural information regarding the full yeast kinetochore comes from electron microscopy (EM; Pesenti et al., 2022; Yatskevich et al., 2022) and fluorescence microscopy studies. EM studies revealed the overall shape of the budding yeast kinetochore (Gonen et al., 2012; McIntosh et al., 2013), but could not assign most proteins in the electron density maps (Hinshaw and Harrison, 2019; Yan et al., 2019). On the other hand, conventional fluorescence microscopy has provided information about the position of several kinetochore components along the spindle axis (Aravamudhan et al., 2014; Haase et al., 2013; Joglekar et al., 2009). However, this approach can only reveal the structural average of all kinetochores, because individual complexes are smaller than the resolution limit of conventional light microscopy (approximately 250 nm (Abbe, 1873)) and clustered. As a result, in budding yeast all 16 kinetochores are observed as one (during interphase) or two fluorescent spots (mitosis; Joglekar et al., 2006), and fine structural details of individual kinetochores cannot be observed. Thus, a comprehensive understanding of the structure of the kinetochore is still missing.

In the budding yeast kinetochore, built on a short centromere sequence (approximately 125 bps; Clarke and Carbon, 1980), the microtubule is captured by multiple copies of the Ndc80c and Dam1c. Precisely how many complexes are present, however, remains controversial, with estimates ranging significantly. To examine this question, previous studies used fluorescence microscopy to quantify the absolute copy numbers of the major kinetochore components. In this approach, the protein of interest was tagged with a suitable fluorescent protein. The brightness of the studied protein was then compared to a reference protein tagged with the same fluorophore (Joglekar et al., 2006, 2008; Lawrimore et al., 2011; Dhatchinamoorthy et al., 2017). These studies generally agreed that the outer kinetochore proteins are the most abundant and the inner kinetochore proteins the least abundant. Ndc80 has been shown to be present in 6 to 19 copies per kinetochore. Smaller or equal amounts were found for the MIND complex (4 to 7 copies) and Spc105 (4 to 5 copies). The COMA complex was shown to be present in 2 to 5 copies. Within the inner kinetochore, Cep3 was found to have 2 to 3.4 copies, Mif2 2 to 3.6 copies, and Cnn1 and Cse4 2 to 6 copies (Shivaraju et al., 2012; Wisniewski et al., 2014). The differences among the results may arise from the choice of the counting reference, cell cycle stage, fluorescent protein, method and optical system used (Joglekar et al., 2008). Such large discrepancies prevent generating a detailed structural model. Open fundamental questions include: How do the Mif2 and Cnn1 assembly pathways quantitatively contribute to the copy number of Ndc80c? How many COMA complexes exist within the budding yeast kinetochore?

Another extensively debated question in the field is the exact stoichiometry at centromeres of the histone protein Cse4. (Clarke and Carbon, 1980; Ng and Carbon, 1987; Keith and Fitzgerald-Hayes, 2000). To date, a series of alternative structures have been proposed to define the nature of the centromeric nucleosome. These hypotheses include hemisome (Bui et al., 2012; Dalal et al., 2007), hexameric (Mizuguchi et al., 2007) or octameric configurations (Camahort et al., 2009), where a single or two copies of Cse4 are present (Black and Cleveland, 2011). With regards to Cse4 copy number, biochemical approaches have reported the presence of a single Cse4 nucleosome at centromeres (Furuyama and Biggins, 2007; Krassovsky et al., 2012). In contrast, *in vivo* studies showed a high variability of Cse4 copy number per kinetochore, ranging from 2 (Dhatchinamoorthy et al., 2017; Shivaraju et al., 2012; Wisniewski et al., 2014) up to 4 - 6 copies (Lawrimore et al., 2011). Interestingly, also the very first SMLM-based counting of Cnp1, the Cse4 homologue in fission yeast, reported 6 - 7 Cnp1 copies per spindle microtubule (Camahort et al., 2009; Lando et al., 2012). Therefore, the identity and copy number of the centromeric nucleosome is still an unanswered question in the centromere and kinetochore fields.

Super-resolution microscopy, and specifically Single-Molecule Localization Microscopy (SMLM; Betzig et al., 2006; Hess et al., 2006; Rust et al., 2006), achieves nanometer resolution combined with molecular specificity, and has the potential to bridge this gap in our knowledge. It has been used to get structural insights into the organization of multi-protein complexes such as the nuclear pore complex (Szymborska et al., 2013), the endocytic machinery (Mund et al., 2018; Sochacki et al., 2017), centrioles (Sieben et al., 2018) or synaptic proteins (Dani et al., 2010). In this study, we use SMLM to determine the location of key proteins and their copy numbers with single kinetochore resolution in *S. cerevisiae* cells (Figure 1). From these data, we built a comprehensive model of how the major components are positioned and what their stoichiometry is in the budding yeast metaphase kinetochore *in situ*.

## Results

### Individual kinetochores can be observed with SMLM

In order to determine whether SMLM can be used to visualize individual kinetochores, we imaged yeast cells in which Ndc80 was endogenously tagged with mMaple, and Spc42 (spindle pole body protein) with GFP (Figure 1B). When we imaged unsynchronized cells, we observed that in interphase cells all kinetochores are packed within a small cluster with a size below the resolution limit of standard microscopy, with the tendency to organize into a rosette-like configuration similar to what is observed in human cells in early prometaphase (Figure 1B; Chaly and Brown, 1988; Jin et al., 2000; Bystricky et al., 2005). In metaphase, kinetochores did not generate a metaphase plate but rather organized into two sister kinetochore clusters (Figure 1B). In late mitosis, the separation of the sister kinetochore clusters increases (Figure 1B; Joglekar et al., 2006). At this late stage of division, their high density did not allow us to resolve individual kinetochores with SMLM. In conclusion, SMLM allows visualizing single kinetochores within the budding yeast spindle in interphase and metaphase.

### Dual-color SMLM quantifies positions of kinetochore proteins along the metaphase spindle axis

In order to resolve structural details of individual kinetochore complexes, we used dual-color super-resolution imaging of two kinetochore proteins along the spindle axis. The distances were measured in a single dimension, with a possible tilt of the spindle axis introducing only a minimal error (maximum error = 6.3%, mean error = 2.1%; see Figure S1 and Methods). We focused on essential kinetochore components and included proteins that have been mapped with diffraction-limited microscopy (Joglekar et al., 2009), for which we could improve the positioning accuracy, and proteins that have never been visualized previously. Unless indicated otherwise, we used Spc105, labeled with SNAP-tag and the organic dye AF647, as a super-resolved spatial reference to position all other proteins, labeled with mMaple, on the spindle axis. To this end, we analyzed each kinetochore cluster individually by reconstructing superresolution images for the reference and target protein and by determining their relative shifts by image cross-correlation (Figure 1C, Figure S2, and Methods). We only analyzed metaphase cells where both kinetochore clusters allowed for high-quality position measurements. As the two kinetochore clusters have an opposite orientation on the spindle axis, minor registration inaccuracies between the channels share the same amount but opposite signs, therefore cancelling each other out (Figure S2D). This allowed us to determine the pairwise distances between 15 pairs of kinetochore proteins, all labeled at their C-termini (Figure 2). We further validated this approach with an independent analysis, in which we directly measured the distance of the proteins in individual kinetochores (Figure S3) and obtained highly similar results. Our measurements of different kinetochore proteins were internally consistent, as the sum of the measured Ndc80 - Spc105 (13.6 ± 1.2 nm; mean ± SEM) and Spc105 - Ctf19 (14.9 ± 1.7 nm) distances is close to the measured Ndc80 - Ctf19 distance (24.9 ± 1.8 nm; Figure 2 inset). These data agree reasonably well with previous diffraction-limited dual-color microscopy studies with noticeable exception of positions of MIND components (for comparison, see Table S1 and Figure S4; Joglekar et al., 2009). Furthermore, we found that the C-termini of Ndc80 and Nuf2 are in close proximity with a distance of 3.3 nm ± 1.5 nm (Figure 2), which agrees well with a distance of 3.6 nm, as determined from a crystal structure (Valverde et al., 2016), adding another validation. In summary, these data show that SMLM dual-color imaging is suitable to measure intra-kinetochore protein distances in budding yeast.

**Figure 2.**
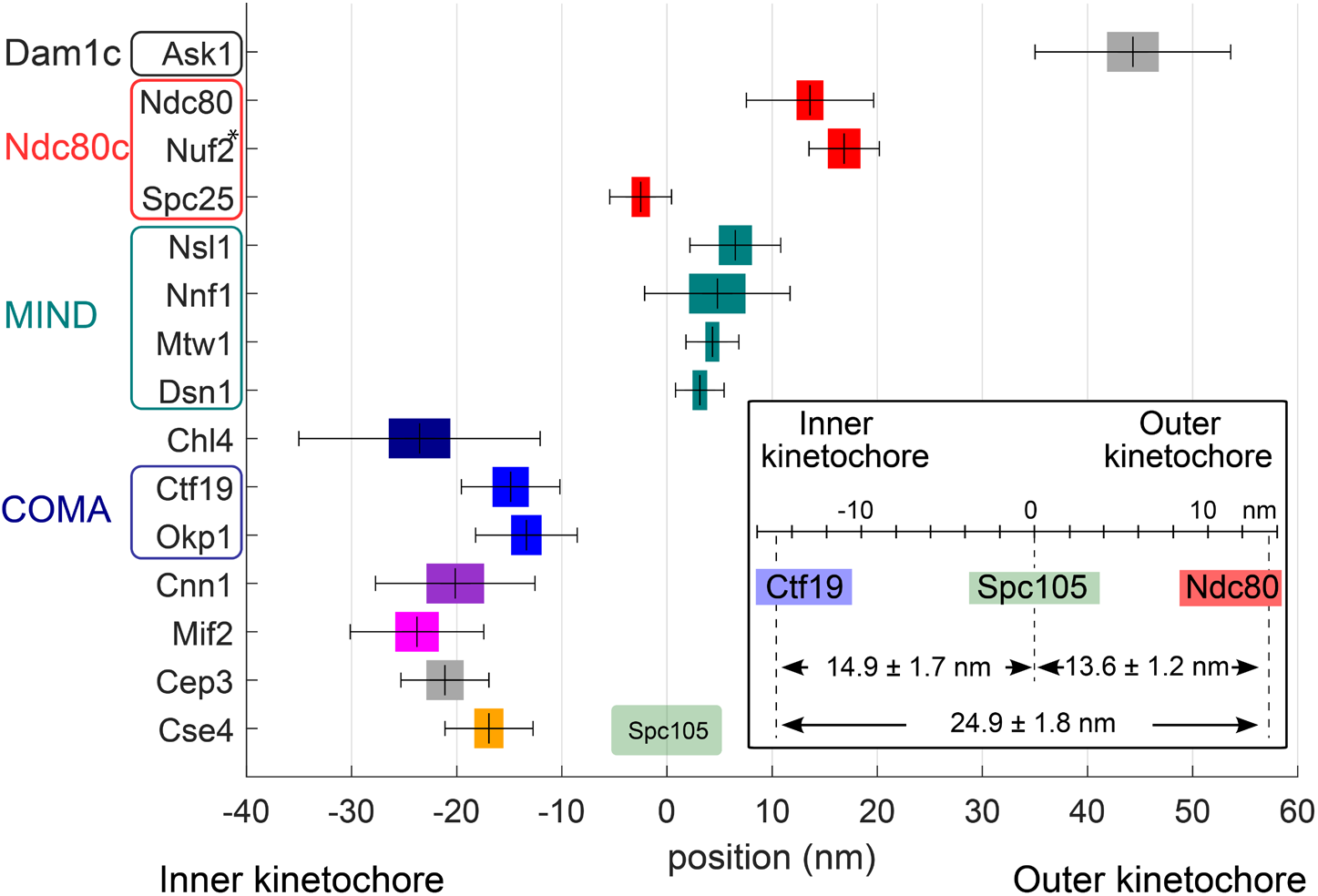
Position of 15 kinetochore proteins along the spindle axis with Spc105 as a reference point. All proteins were tagged at their C-termini. The mean distance is plotted with the standard error of mean (SEM, colored box) and standard deviations (SD, whiskers). The inset depicts control measurements showing consistency in pairwise distance measurements ± SEM among three proteins. See Table 1 for values. *The position of Nuf2 is based on the measured pair Ndc80-Nuf2.

We found the C-termini of Cse4 and Cep3 to be positioned within 1.5 nm away from each other at the centromeric site. Also, Mif2 and Cnn1 cluster together, which is consistent with their function within the inner kinetochore (Figure 2) but are around 3 nm away from the Cse4, towards the Cep3 site. Interestingly, we measured the position of Chl4 to be only 0.3 nm away from Mif2, but more distant from the COMA complex (8.9 nm). We find that Ctf19 and Okp1 (COMA components) are −14.9 ± 1.7 nm and - 13.4 ± 1.4 nm away from Spc105, respectively, towards the centromere (Figure 2).

Next, we found that Nnf1, Nsl1, Mtw1, and Dsn1, which all belong to the MIND complex, are between 3.1 nm and 6.5 nm away from Spc105 in the outward direction (towards the microtubule). This is consistent with a crystal structure of MIND in yeast and human and with the known binding site of the KNL1^Spc105^ C-terminus on the MIND complex (Dimitrova et al., 2016; Hornung et al., 2014; Kudalkar et al., 2015; Petrovic et al., 2016; Petrovic et al. 2014). While the C-terminus of Spc25 is adjacent to the C-termini of both Spc105 and MIND (Figure 2), the C-terminus of Ndc80 occupies a more outward position. Finally, the Ask1 subunit of Dam1c is positioned around 40 nm away from Spc105 in the microtubule direction.

Our data also contain information about the distribution widths of kinetochore proteins perpendicular to the spindle axis. We extracted this information using auto-correlation analysis. We found that the width of the distribution correlates to the position of the protein along the spindle axis (Figure S5). Using auto-correlation of simulated ring distributions with different radii as references, we found that most inner kinetochore proteins are distributed within a radius of 10 to 15 nm of the kinetochore center and most outer kinetochore proteins within a radius of ~15 nm. The wider distributions of the outer kinetochore proteins can be explained by the presence of microtubule, which has a radius of ~12.5 nm, occupying the central space.

### Counting kinetochore protein copy numbers with quantitative SMLM

In order to estimate the protein copy numbers of the major kinetochore components, we used a quantitation approach based on reference standards for super-resolution counting (Thevathasan et al., 2019). Here, the target complex is imaged under identical conditions as the reference standard, tagged with the same fluorophore (mMaple). The copy number of the unknown complex can be directly calculated from the known copy number of the counting standard and the relative number of detected localizations. We selected Nup188, a protein component of the nuclear pore complex (NPC), as a bright and easy to segment counting reference complex (Figure 1C; Thevathasan et al., 2019). Nup188 has 16 copies per budding yeast NPC (Kim et al., 2018). We mixed the reference strains containing Nup188-mMaple and Abp1-GFP as an identification marker with the target strains containing mMaple-labeled kinetochore proteins and imaged them on the same coverslip to ensure identical imaging conditions. We further improved the accuracy by employing highly homogenous illumination (Deschamps et al., 2016) throughout the entire field of view. We usually acquired images of 600 NPCs and 200 kinetochore clusters per experiment. We only analyzed kinetochore clusters that were close to the focal plane to ensure that the analyzed kinetochore proteins did not exceed the imaging depth (see Figure S1, Figure S6 and Methods). This allowed us to precisely calculate the copy numbers of kinetochore proteins.

For the inner kinetochore, we first quantified Cse4. Previous reports have indicated that only internal genetically-encoded fluorescent tagging of Cse4 is compatible with its physiological function, while Nor C-terminal tagging renders cells less viable (Wisniewski et al., 2014). However, in our experiments, we found that both internal and C-terminal tagging of Cse4 were compatible with viability. Furthermore, our counts were essentially identical, with 4.2 ± 2.0 (standard deviation; SD) copies of the histone Cse4 when it is tagged internally and 4.8 ± 2.4 (Figure 3) when the tag is localized at its C-terminus. Cep3 was found in 4.2 ± 2.1 copies. Mif2 and Cnn1 are present in 3.5 ± 1.7 and 2.1 ± 1.3 copies/kinetochore, respectively. The COMA complex component Ctf19 has 4.1 ± 1.9 copies, and the COMA and Mif2 binder Chl4 is present in 1.8 ±1.0 copies. The outer kinetochore proteins are present in higher copy numbers: 7.6 ± 3.4 copies of Spc105, 7.2 ± 3.2 of Dsn1, 10.9 ± 5.0 of Ndc80 and 24.9 ± 11.0 of Ask1 (Figure 3).

**Figure 3.**
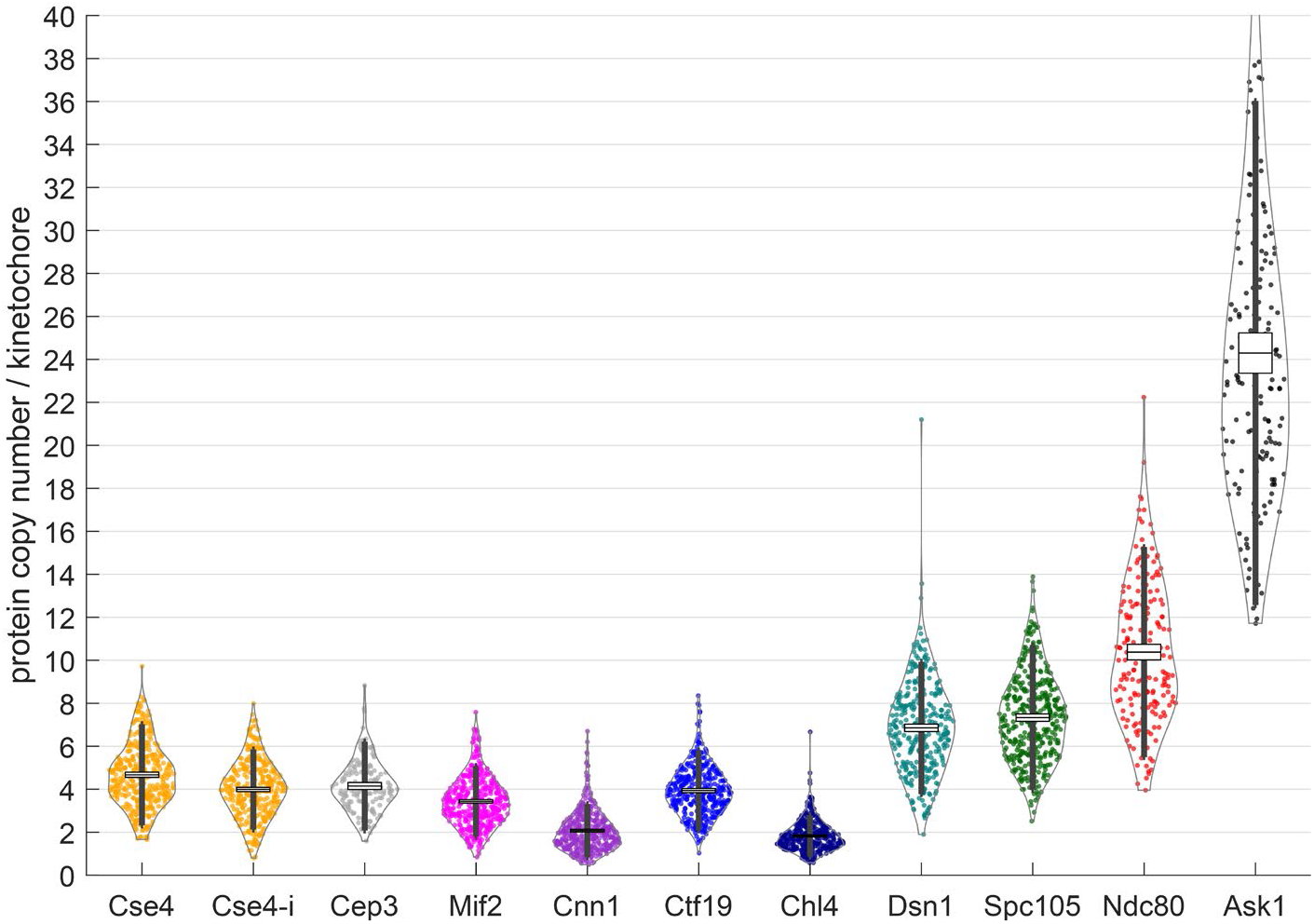
Protein copy numbers per kinetochore. measured with Nup188-mMaple as a counting reference standard. Each data point corresponds to one kinetochore cluster. All proteins were tagged at their C-termini, except Cse4-i that was tagged internally. Boxes denote average copy numbers and standard error of means, and whiskers denote standard deviations. For each protein, two independent experiments were performed and pooled (see Methods for details).

Different lifetimes of the Nup188 and kinetochore proteins could lead to different maturation efficiencies of the mMaple tag and consequently to systematic errors in the counting measurements. To investigate the effect of tag maturation, we transiently stopped protein translation with 250 μg/ml cycloheximide (CHX) and performed our counting measurements one hour after this treatment (Figure S3). Although we observed minor changes in copy numbers, the overall effect of CHX was small. The noticeable exception was internally tagged Cse4 for which 30 – 40% reduction of the signal was seen. We conclude that tag maturation does not grossly affect our measurements of protein copy number. We have not noticed any growth defects that may have arisen from the tagging in our experiments, but we do not exclude a possibility of minor effects. However, our data is consistent with the previous measurements suggesting that our C-terminal tagging did not introduce any artefacts (Joglekar et al., 2006, 2008, 2009; Lawrimore et al., 2011; Pekgöz Altunkaya et al., 2016; Dhatchinamoorthy et al., 2017).

### Quantitative model of the budding yeast kinetochore

We then integrated all protein copy numbers (Figure 3) and protein-protein distance measurements along the spindle axis (Figure 2) in a model of the structural organization of the budding yeast kinetochore (Figure 4). Based on their close proximity (Figure 2), their known tendency to dimerize (Cohen et al., 2008) and non-centromeric DNA interactions we positioned at the centromeric site two copies of Cse4, a dimeric CBF3 subunit (with two Cep3 dimers), Mif2 dimer and two copies of Cnn1. Roles of the additional copies of Cse4, Mif2, CBF3 and COMA molecules detected by our measurements (indicated in Figure 4 by dashed lines) needs to be further investigated. In addition, we only included essential structural information (protein structure and binding partners) well established in the field. Specifically, we did not divide the inner kinetochore components by their centromeric-proximal, peri-centromeric or other nuclear localization. Next, we placed all C-termini of MIND proteins away from COMA. We then positioned seven copies of Spc105 and of MIND and ten globular Spc25-containing ends of Ndc80c in close proximity to each other. Four unbound Ndc80c were left for Cnn1 binding. Finally, we present Dam1 complexes as an oligomeric structure surrounding the microtubule.

**Figure 4.**
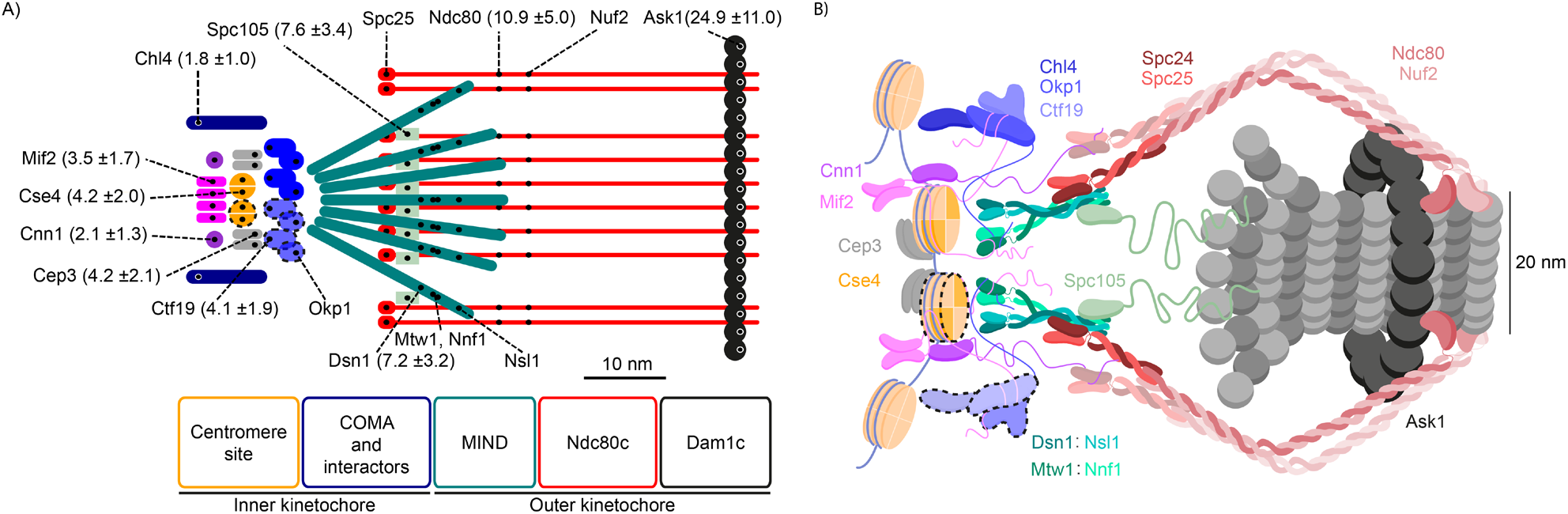
Structural model of the budding yeast kinetochore. **A. Quantitative schematic model** based on the position and protein copy numbers measured with SMLM. The position of the label is shown as a small black dot. Values in the brackets are the estimates of the number of proteins per kinetochore +/- SD. **B. Illustrative structural model** that we built by integrating our position and copy number measurements with previous models (Jenni et al., 2017; Fischböck-Halwachs et al., 2019; Hamilton et al., 2019; Ustinov et al., 2020). Dashed lines indicate potentially accessory (non-centromeric) copies (see Discussion for details). For simplicity, only two copies of COMA, MIND and Spc105 and four copies of Ndc80c are shown in **B**.

## Discussion

In this study, we used single-molecule localization microscopy to position 15 kinetochore proteins along the spindle axis length in metaphase and measured the copy numbers of 10 representative kinetochore components (Figure 4), giving new insights into the structural organization of the budding yeast kinetochore *in vivo*.

### Kinetochore subunits are organized functionally along the spindle axis

Using dual-color SMLM, we mapped the relative positions of 15 kinetochore proteins along the spindle axis with nanometer precision. The resulting position map clearly showed that the structural organization of kinetochore proteins correlated with their function and confirmed the general structure of the inner and the outer kinetochore. Kinetochore proteins known to interact with each other were found in close proximity in our analysis, validating their interactions and our approach.

Within the centromere-proximal region, which is more than 20 nm away from the outer kinetochore and the reference protein Spc105, Cse4 and CBF3 (measured with its constituent Cep3) colocalize with each other as well as with the C-termini of both outer kinetochore receptors Mif2 and Cnn1. The Cep3 dimer, within the CBF3 complex, binds CDEIII DNA and participates in Cse4-containing centromere deposition (Leber et al., 2018; Yan et al., 2018; Zhang et al., 2018; Hinshaw and Harrison, 2019). Cnn1 does not seem to bind the centromeric nucleosome directly but its localization depends on Mif2 (Schmitzberger et al., 2017). These generate the base for further kinetochore assembly. Additionally, we find Chl4 within the centromere-proximal region as well, which is in line with Chl4 interacting with Mif2, the Cse4-containing nucleosome and, electrostatically, with DNA (McKinley et al., 2015; Pentakota et al., 2017). The COMA complex (as measured with Ctf19 and Okp1) occupies the intermediate position, 15 to 20 nm from Spc105, bridging the inner with the outer kinetochore (Hinshaw and Harrison, 2019; Hornung et al., 2014). The outer kinetochore components (Spc105, MIND, Ndc80c, Dam1c) are more distal from the centromere and create the microtubule-interacting module, with the Ndc80c and Dam1c directly binding the microtubule surface (Cheeseman et al., 2006; Ciferri et al., 2008; Wei et al., 2007). All C-termini of the MIND complex are localized more than 10 nm away from COMA, suggesting that all N-terminal regions of MIND proteins lie relatively close to the complex. This is supported by numerous previous biochemical and optical studies (Aravamudhan et al., 2014; Dimitrova et al., 2016; Petrovic et al., 2016). The distance between the position of COMA and the C-termini of MIND implies a possible tilt between the longer axis of MIND and spindle as the total length of MTW1 is around 20 nm (Hornung et al., 2011). The structured segment of Spc105, the reference point, is positioned close to the C-termini of MIND, as was proposed previously using structural approaches (Petrovic et al., 2014). The Ndc80c is an elongated heterotetramer. The C-termini of two of its constituents (Spc25 and Ndc80) are 14.1 nm away from each other, a few nanometers less than the maximum length of this region observed in the purified sample (Wei et al., 2005; Valverde et al., 2016). The discrepancy between structural data of MIND and Ndc80c in our measurements can be explained as an existing tilt of both complex to the spindle axis. The tilt may have an implication in response to a tension during bi-orientation and in accommodation of Ndc80c binding to a microtubule surface. Based on the distance between Okp1 and Ndc80 the tilt can be estimated to be around 46 degrees. Finally, another complex assembles around the positive end of a spindle microtubule—Dam1c, placed some 40 nm outward from Spc105. As its maximum outer diameter is around 50 nm (Ramey et al., 2011), Ndc80c must overcome this barrier in order to reach the microtubule surface.

Generally, our results align with previous biochemical complex reconstitutions, protein interaction studies and with the majority of optics-based distance measurements. Compared to previous optical measurements, the tenfold higher resolution in our study greatly improved the accuracy of position estimates with the single-kinetochore resolution removing a bias from proteins that are not incorporated in kinetochores but nonspecifically enriched in the spindle region. Thus, we found the C-termini of the MIND complex positioned within the outer kinetochore region between Spc105 and the Ndc80c. Here, the C-terminus of Dsn1 highly overlap with Spc105 position, whereas Nnf1, Mtw1, Nsl1 C-termini extend towards the position of Ndc80. This adjusts a previous study that measured the distance between the diffraction limited spots of fluorescently-tagged kinetochore proteins in living cells and found the C-termini of Mtw1, Nsl1 and Dsn1 7 nm away from Spc105 in the direction of the centromere, whereas Nnf1 was shown to fully colocalize with Spc105 (Joglekar et al., 2009; Table S1 and Figure S4).

Our results on COMA and Ndc80c are also compatible with previous studies, but we add position information about important proteins that have not yet been mapped, namely the Cse4 C-terminus, Cef3, Mif2, Cnn1 and Chl4.

### Copy numbers of the major kinetochore components

The quantitative SMLM counting approach recently developed in our lab (Thevathasan et al., 2019) allowed us to precisely measure the copy number of specific proteins per kinetochore (Figure 3). One highly debated question in the field is the composition of the centromeric nucleosome and, with this, the copy number of Cse4 within individual kinetochores. There is a strong disagreement between biochemical and in-situ assays. Using chromatin immunoprecipitation (ChiP), only a single centromere-specific nucleosome can be recovered (two Cse4 copies; Furuyama and Biggins, 2007; Krassovsky et al., 2012; Pekgöz Altunkaya et al., 2016), which is also supported by a disc-like shape structure of the nucleosome observed by electron microscopy within a yeast metaphase spindle (McIntosh et al., 2013). On the other hand, microscopy data point to higher copy numbers of Cse4, exceeding the expected single centromere-specific nucleosome per kinetochore (Lawrimore et al., 2011). ChiP methods may not be able to detect the additional Cse4 due to their limit of detection (Lawrimore et al., 2011).

In our study, we find up to four copies of Cse4 per kinetochore (Figure 3), independently of whether tagging was internal (near the N-terminus) or at the C-terminus (Wisniewski et al., 2014) though the decrease of a copy number was observed upon cycloheximide treatment (Figure S7). To obtain further information about the centromere environment, we measured the copy numbers of the Cse4-binders Mif2 and Cep3 (CBF3 complex). We found that Cep3 have an equal copy number of 4 per kinetochore and Mif2 may be present as two dimers (four copies). The CBF3 complex containing two Cep3 dimers was shown to potentially allocate to a kinetochore (Yan et al., 2018). However, Cep3 exhibits also non-kinetochore localization (Joglekar et al., 2006). It is worth noting that in other organisms the CENP-C dimer may interact with two centromeric nucleosomes distinguishing the budding yeast centromere even more (Carroll et al., 2010; Guse et al., 2011; Watanabe et al., 2019; Ali-Ahmad et al., 2019; Walstein et al., 2021). Our study supports the notion that, among other inner kinetochore components, non-centromeric Cse4 may play a role in maintaining the “point” centromere by serving as a spare module (as discussed in Scott and Bloom, 2014).

In our study, we found four copies of Ctf19 but only two copies of Chl4 per kinetochore. Structural studies have shown only two COMA complexes within a kinetochore (Hinshaw and Harrison, 2019). Thus, we placed additional COMA copies as accessory (non-centromeric; Figure 4). It is widely accepted that N-termini of both Mif2 protein and COMA subunits allow and regulate assembly of the outer kinetochore module (Dimitrova et al., 2016; Petrovic et al., 2016; Przewloka et al., 2011; Screpanti et al., 2011). With a total of two interaction sites from a Mif2 dimer and two COMA, a budding yeast kinetochore may build up to four copies of MIND. This would leave additional copies unbound. However, crystallographic packing of MIND reveals potential oligomerization (Dimitrova et al., 2016) allowing us to place all complexes within the kinetochore. This in turn would bring equal or similar amount of Spc105 and Ndc80 complexes (Petrovic et al., 2014). Indeed, we observed 6 - 8 MIND complexes and an equal number of Spc105. Consistently with others (Joglekar et al., 2006; Dhatchinamoorthy et al., 2017) we found more Ndc80 than Spc105 or MIND per kinetochore. However, the ratio between estimated copy numbers of Cse4 and Ndc80 in the current analysis is 2.5. Thus, it is different from in the aforementioned studies where Ndc80 is 4 times more abundant. The additional 2 Ndc80 can be bound by the last outer kinetochore receptor Cnn1. In regional kinetochores, CENP-T, the Cnn1 orthologue, recruits up to three Ndc80c to the outer kinetochore (Huis in ’t Veld et al., 2016). In budding yeast, each Cnn1 can bind two Ndc80c (Pekgöz Altunkaya et al., 2016). The binding is regulated by Cdk1- and Mps1-dependent phosphorylation of Cnn1 (Malvezzi et al., 2013). The decreasing activity of the aforementioned kinases may allow the Cnn1-Ndc80 interaction to be more permissive. Our observations were limited only to metaphase. Therefore, the results are consistent with one Cnn1 binding to a total of two to three Ndc80 per kinetochore. Yet, when Ndc80c copy numbers are estimated in Cnn1-deleted strains the copy number is not altered (Pekgöz Altunkaya et al., 2016; Dhatchinamoorthy et al., 2017) or the change may be minimal when MIND-Ndc80c binding pathway is impaired (Lang et al., 2018). This points to the redundancy of Cnn1 in budding yeast when the mitotic checkpoint is not compromised, or to a dynamic nature of the Ndc80-Cnn1 interaction.

We have estimated slightly higher copy number of Ask1 protein (a single Ask1 molecule is present in a Dam1c monomer) per kinetochore than an earlier work (16-20 copies; Joglekar et al., 2006). In general, 17 copies form a complete microtubule-encircling Dam1c ring (Ng et al., 2018). However, different configurations of Dam1c oligomerization (one and two partial/complete rings) might exist on one microtubule even in the same cell (Ng et al., 2018). Two Dam1c rings on each microtubule have also been suggested (Kim et al., 2017). These altogether may explain the variation and higher mean copy number of Ask1 we quantified.

### Summary

Taken together, we employed the high resolution of SMLM to substantially improve the accuracy of previous stoichiometry and intra-kinetochore distance estimates and obtained a comprehensive model of the structural organization of the kinetochore in budding yeast *in situ* (Figure 4), revising previous models (Jenni et al., 2017; Fischböck-Halwachs et al., 2019; Hamilton et al., 2019; Ustinov et al., 2020). This model adds additional valuable information to understand how the metaphase kinetochore is structurally organized *in situ* by overcoming the resolution limit present in the previous studies.

In an independent investigation, a similar methodology was used to assess protein composition and distances of *S. pombe* kinetochores (Virant et al., 2021). Their results are in excellent agreement with ours, as expected from the high conservation of kinetochore components across the two yeast species (Hooff et al., 2017), validating our respective approaches. One main difference is the Cse4:COMA ratio, which is 1:0.9 in budding yeast and 1:2.1 in fission yeast, pointing to intrinsic stoichiometry changes between point and regional kinetochores. In conclusions, our quantitative SMLM methods provide a strong basis for future studies, for instance how kinetochore components are organized perpendicular to the spindle axis and how this relates to the kinetochore-microtubule binding management, how structure and stoichiometry change throughout the cell cycle or how kinetochores are organized in other organisms. Our methods are not restricted to kinetochores, but will enable quantitative measurements of the stoichiometry and structure of other multi-protein assemblies *in situ*.

## Acknowledgements

We thank Andrea Musacchio and Ulrike Endesfelder for feedback on the manuscript and Katharina Lindner for her work on the imaging of the dual color strains. This work was supported by the European Research Council (grant no. ERC CoG-724489 to JR), the Human Frontier Science Program (RGY0065/2017 to JR) and the European Molecular Biology Laboratory.

## Author contributions

K. C. and J.R. conceived the study. K.C., S.J.H., Y.-L.W. and M.S. performed experiments. K.C., Y.-L.W., L.N., M.S. and J.R. analyzed data. K.C., Y.-L.W. and L.N. visualized the results. J.R. supervised the study. K.C., Y.-L. W., D.C., and J.R. wrote the manuscript with input from all authors.

## Methods

### Yeast strain generation

All strains used in the study (Table S2) were derived from *S. cerevisiae* MKY0100 strain (S288c derivative), a kind gift from the Kaksonen lab (University of Geneva). The strains for endogenous expression of fluorescently-tagged kinetochore proteins were created by homologous recombination using PCR-based C-terminal tagging cassettes (Janke et al., 2004). The cassettes were created by amplification of DNA regions of respective pFA6a plasmids (Mund et al., 2018) encoding mMaple (McEvoy et al., 2012) or SNAPf tag (Sun et al., 2011). The Cse4-mMaple-Cse4 strain was created analogically to Wisniewski et al., 2014. Cse4 and mMaple sequences were amplified by PCR and ligated into pFA6a vector replacing a tag sequence. Subsequently, PCR product encoding Cse4-mMaple-Cse4-HIS3MX6 was used to transform yeast competent cells by standard lithium–acetate protocol. Correct genome integrations in transformed yeast cells were checked by PCR.

### Sample preparation

24 mm round coverslips were cleaned in HCl/Methanol overnight and then rinsed with water. Additionally, the coverslips were cleaned using a plasma cleaner to remove residual organic contaminations. Coverslips were then coated with 15 μl of Concanavalin A (4 mg/ml in PBS; Sigma C2010), dried overnight at 37°C, and before use rinsed with water to remove residual PBS. The coverslip was covered with ~100 μl of a cell suspension and incubated for 15 min.

For mMaple imaging, 2 ml of yeast logarithmic culture was grown in SC-Trp, spun down (2500 rpm, 3 min) and resuspended in 100 μl of the medium. In case of the control experiments with cycloheximide treatment, 250 μg/ml of cycloheximide (in DMSO) was added to cells 1 hr before immobilization. Cells immobilized on Concanavalin A-coated coverslips were fixed in 4% paraformaldehyde, 2% sucrose in PBS for 15 min at room temperature. Fixation was quenched by 2 washes in 100 mM ammonium chloride, pH 7,5 in PBS for 20 min. Finally, the sample was rinsed with PBS several times. The coverslip was mounted on a microscope stage and covered with 50 mM Tris-HCl, pH 8 in 95% D_2_O.

For single- and dual-color imaging with SNAP, the cells were immobilized, fixed and washed the same way. Subsequently, the cells were permeabilized by 0.01% digitonin in 1% BSA solution for 30 min at room temperature under moist conditions. The sample was then washed in PBS. The sample was labeled with 1 μM SNAP-Surface Alexa Fluor 647 in 1% BSA solution for 2 h at room temperature under moist conditions. Finally, the sample was washed in PBS 3×5 min. The sample was mounted in a microscope stage and covered with the blinking buffer consisting of 50 mM Tris-HCl, pH 8, 10 mM NaCl, 10% (w/v) D-glucose, 500 μg/ml Glucose oxidase, 40 μg/ml Catalase in 90% D_2_O (Thevathasan et al., 2019). The blinking buffer for Alexa Fluor single-color or dual-color imaging was supplemented with 35 mM or 15 mM MEA (mercaptoethylamine), respectively.

### Microscopy

The SMLM acquisitions were performed with the two custom-build microscopes, analogically as in (Mund et al., 2018), and with custom-developed EMU interface (Deschamps and Ries, 2020). Microscope 1 was used for low-throughput single- and dual-color imaging. Before dual-color experiments, a bead calibration with 100 nm Tetra-Speck beads for a faithful channel overlay was performed. The splitting of the emission signals was achieved with a 640 nm long pass dichroic mirror. The signal from the 640 nm or 562 nm laser excitation was collected through 676/37 nm or 600/60 nm emission bandpass filters, respectively. SMLM measurements were performed with 30 ms exposure time. The UV laser was adjusted automatically to keep the density of localizations constant (Mund et al., 2018). The cells with similarly bright sister kinetochore signals were chosen for each acquisition. Initially, we imaged the cells with the 640 nm laser until the localization density was sufficiently reduced. Then the 561 nm laser was switched on. Typically, we acquired 60000 frames and obtained ~35 nm localization precision for mMaple and 20 nm for Alexa Fluor 647.

Microscope 2 was primarily used for high-throughput single-color mMaple imaging. As in microscope 1, microscope 2 has several available channels - UV (405 nm), green (488 nm laser, 525/50 nm emission bandpass filter), orange (561 nm laser, 600/60 nm emission bandpass filter), red (640 nm - excitation and booster laser, 700/100 nm emission bandpass filter). A focus lock system based on a totally reflected IR laser beam was used to keep the focus constant. In order to keep the illumination of the entire field of view uniform we used homogenous and speckle-free illumination (Deschamps et al., 2016).

For protein counting experiments, two strains expressing the Nup188-mMaple standard and the target kinetochore protein labeled with mMaple were mixed and imaged simultaneously. 225 regions were imaged per coverslip, separated by at least 150 μm to avoid premature mMaple activation. Every acquisition was performed with approximately 100 mW of the 561 nm laser, 25 ms exposure time and the UV laser adjusted automatically to result in a constant, but low density of activated fluorophores. All measurements were performed until all mMaple fluorophores had been activated and bleached. A snapshot of Ndc80-GFP (for kinetochores) or Abp1-GFP (for Nup188-mMaple strain) was automatically acquired, as well as a back focal plane image to exclude acquisitions with air bubbles.

### Single-molecule localization

We used SMAP program package (Ries, 2020) for all data analysis. For single-molecule fitting, candidate localizations were detected by smoothing with a Difference of Gaussians filter and thresholding. Then, the signal was localized by fitting a Gaussian function with a homogeneous photon background, treating the size of the Gaussian as a free fitting parameter. Fluorophores spanning consecutive frames and thus likely stemming from the same fluorophore were merged (grouped) into a single localization. For experiments longer than 5000 frames, cross-correlation based sample drift correction was applied as described in (Mund et al., 2018). Super-resolution images were reconstructed by rendering each localization as a Gaussian with a size proportional to the localization precision. Finally, localizations were filtered by localization precisions to exclude dim emitters and by PSF sizes to exclude out-of-focus fluorophores. If the localization density in the first frames was above the single molecule regime, these frames were discarded.

Dual-color bead images were fitted as described above and used to calculate a projective transformation between the channels.

For high-throughput data we extracted additional parameters for quality control such as the number of localizations and the median localization precision, photon count, PSF size and background, and used them in combination with the BFP images to exclude poor measurements that resulted from air bubbles in the immersion oil or acidification of the buffer.

### Z-position bead calibration

The preparation of the bead sample is similar to the 3D bead calibration described in (Thevathasan et al., 2019). Briefly, Tetra-Speck beads (0.75 μL; catalog no. T7279, Thermo Fisher) were diluted in 360 μL H2O, mixed with 40 μL 1 M MgCl_2_ and put on a coverslip in a custom-manufactured sample holder. After 10 min, the mix was replaced with 400 μL H_2_O. Using Micro-Manager (Edelstein et al., 2014), about 20 positions on the coverslip were defined and the beads were imaged acquiring *z* stacks (−1 to 1 μm, 10 nm step size) using the same filters as above. Images of beads were then localized to quantify their PSF sizes. Based on the PSF sizes and the stack positions, the z positions of fluorophores can be calibrated (Figure S1D).

### Quantification of distances between kinetochore proteins

We quantified distances between kinetochore proteins based on a cross-correlation analysis. Before the analysis, in a dual-color SMLM data set, localizations with localization precision > 20 nm for Alexa Fluor 647 and > 25 nm for mMaple channels or PSF size <100 nm or >160 nm were removed. Only the in-focus structures (mean PSF size ≤ 135 nm) were kept for the analysis. One color/channel (usually the channel of Spc105 unless specified otherwise) was defined as the reference, and the other as the target. We started by manually collecting kinetochore clusters (sites) and grouped both kinetochore clusters of the same mitotic spindle as a pair (Figure S2A). For each pair, a line was manually drawn to represent the spindle axis, which the kinetochore clusters distributed along. Next, to take the opposite direction of chromosomes pulling by each kinetochore cluster of the pair into account, the axial direction was defined as pointing towards the center of the spindle (Figure S2A). As shown in Figure S2B, each kinetochore cluster/pair of kinetochore clusters went through the same analysis steps (Figure S2C and D) for quantifying the distance. First, we calculated the image cross-correlation between two reconstructed super-resolution images corresponding to the two channels for each kinetochore cluster separately. From the maximum position of the cross-correlation map we determined the average distance between the two proteins along the spindle axis. To exclude that residual transformation errors caused e.g., by chromatic aberrations, we always analyzed the two paired kinetochore clusters together. Due to their close proximity, we expect similar registration errors, which cancel out when calculating the average protein distance because of the opposite orientation of the kinetochore clusters. As a result, each spindle resulted in one average distance value. Using Spc105 as a reference in most data sets, we could position all measured proteins along the spindle axis. The number of experiments per kinetochore protein is summarized in Table 1 and Table S3.

**Table 1.** Statistics of kinetochore protein positions along the spindle axis. *The position of Nuf2 is based on the measured pair Ndc80-Nuf2. SD: standard deviation, SEM: standard error of the mean, N: number of spindles.

| Protein | Distance to Spc105 (nm) | SD | SEM | N |
| --- | --- | --- | --- | --- |
| Ask1 | 44.3 | 9.3 | 2.4 | 15 |
| Ndc80 | 13.6 | 6.1 | 1.2 | 25 |
| Nuf2* | 16.9 | 3.3 | 1.5 | 5 |
| Spc25 | -2.5 | 2.9 | 0.8 | 13 |
| Nsl1 | 6.5 | 4.3 | 1.5 | 8 |
| Nnf1 | 4.8 | 6.9 | 2.6 | 7 |
| Mtw1 | 4.3 | 2.5 | 0.6 | 17 |
| Dsn1 | 3.1 | 2.3 | 0.6 | 13 |
| Chl4 | -23.5 | 11.5 | 2.9 | 16 |
| Ctf19 | -14.9 | 4.7 | 1.7 | 8 |
| Okp1 | -13.4 | 4.8 | 1.4 | 12 |
| Cnn1 | -20.1 | 7.6 | 2.7 | 8 |
| Mif2 | -23.8 | 6.3 | 2.0 | 10 |
| Cep3 | -21.1 | 4.2 | 1.7 | 6 |
| Cse4 | -16.9 | 4.2 | 1.3 | 10 |
| Ctf19-Ndc80 | -24.9 | 8.5 | 1.8 | 23 |

### Estimation of the error introduced by axial tilts of spindle axes

We first quantified the average width of kinetochore clusters based on a cylindrical distribution. Specifically, the 1D profile along the diameter of a cylinder convolved with a Gaussian function (σ defined as the mean localization precision) was calculated. Such a profile was fitted to kinetochore clusters with the radius as a free parameter.

We localized emitters in the bead z-stacks acquired as described above to obtain their PSF sizes. We then fitted a quadradic curve to the scatter plot of the PSF sizes and z positions of beads. The fitted calibration curve describes the relation between z positions of localizations and PSF size.

The 1D profile of cylindrical distribution with the radius defined as the quantified average width of kinetochore clusters was plugged into the calibration curve to obtain a new calibration curve describing the relation between z position of a kinetochore cluster and its mean PSF size. We then drew a line at mean PSF size = 135 nm, which is the maximal possible value of the analyzed kinetochore clusters (Figure S1E). The maximal axial distance between kinetochore clusters in the same pair 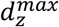 is defined as the distance between the cross-points of the line and the calibration curve. The distance between the two kinetochore clusters in 3D was estimated as 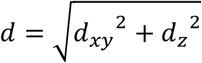, where *d_xy_* is the lateral distance between the two kinetochore clusters. The relative error introduced by the axial tilt is calculated as ε(θ) = (*d* – *d_xy_*)/*d*, where *θ* = cos^-1^(*d_xy_*/*d*) is the tilt angle. The maximum tilt angle *θ^max^* was estimated based on mean lateral distance 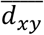 and the estimated maximum axial distance 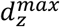. The mean error is then estimated as 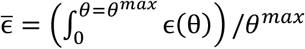.

### Estimations of the widths of kinetochore protein distributions

We used auto-correlation analysis to quantify the widths of kinetochore protein distributions. For each kinetochore cluster, we generated a 2D auto-correlation map. For each map, the auto-correlation values at shifts along the spindle axis < 25 were summed per shift perpendicular to the spindle axis to yield the profile across the shifts. The high auto-correlation value at the shift = 0 was substituted by the value of its neighboring shift. The profile was then normalized to have the maximum of 1 before averaging over all kinetochore clusters of the same kinetochore proteins. To separate the real auto-correlation from its background, two Gaussian functions with a linked parameter μ (position) were then fitted to the averaged profile. The function with the larger fitted parameter σ was considered as the background and then subtracted from the averaged profile. This profile for each analyzed protein is shown Figure S5.

We performed simulations to obtain reference auto-correlation profiles of ring distributions with different radii. Specifically, the 1D profile along the diameter of a ring was calculated per specified radius. To take the experimental localization precision into account, we acquired its binned distribution based on the mMaple channel over all the dual-color data sets. We then convolved the 1D profile with a Gaussian function (σ taken from the bin value) per bin. We then summed the profiles weighed by the frequency of the corresponding bins to form the final profiles. For each final profile, its auto-correlation was then calculated and is shown in Figure S5.

### Protein copy number estimations

To differentiate the yeast strains on the same coverslip, proteins with different cellular distributions were tagged with mEGFP in the reference and target strains (Abp1 for the reference and a kinetochore protein for the target). The GFP signal was checked in the diffraction-limited channel. We then manually segmented the single structures of the reference (NPCs) and the target (kinetochore clusters) in respective strains. Before further analysis, localizations with localization precision > 15 nm or PSF size <100 nm or >170 nm were removed. Only the in-focus structures (mean PSF size ≤ 135 nm) were retained in the analysis. For the reference, NPCs at the edge of the nucleus or too close to neighboring structures were excluded. We then determined the number of localizations in a circular ROI of a diameter of 150 nm. For a target structure, we only picked kinetochore clusters that have two foci in the GFP channel to ensure metaphase kinetochore clusters. We then determined the number of localizations in the manually-created polygon enclosing the kinetochore cluster. When paired kinetochore clusters were too close to each other, they were segmented as one entity and its localizations were divided by 2. The copy number calibration factor for each dataset was calculated as *F_n_* = *L_n_/N_n_*, based on the stoichiometry of Nup188 (Table S4). Here *L_n_* is the mean quantified localizations per NPC and *N_n_*=16 is the known copy number of Nup188 per NPC. Then the copy number *N_k_* of a target protein per kinetochore was calculated as 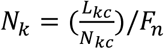, where *N_kc_* = 16 is the number of kinetochores per kinetochore cluster and *L_kc_* is the mean quantified localizations per kinetochore cluster. To take the variation of the NPC localizations into account, the standard deviation of the kinetochore protein copy number was 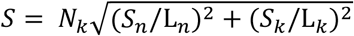, where *S_k_* and *S_n_* are the standard deviations of the localizations for NPC and kinetochore protein *L_k_* and *L_n_* are the respective sample sizes. Finally, the pooled copy number and standard deviation of replicates were 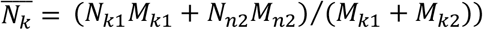 and 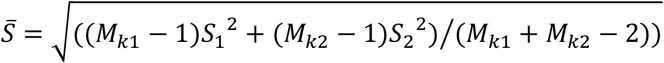, respectively.

## Supplementary Figures

**Figure S1.**
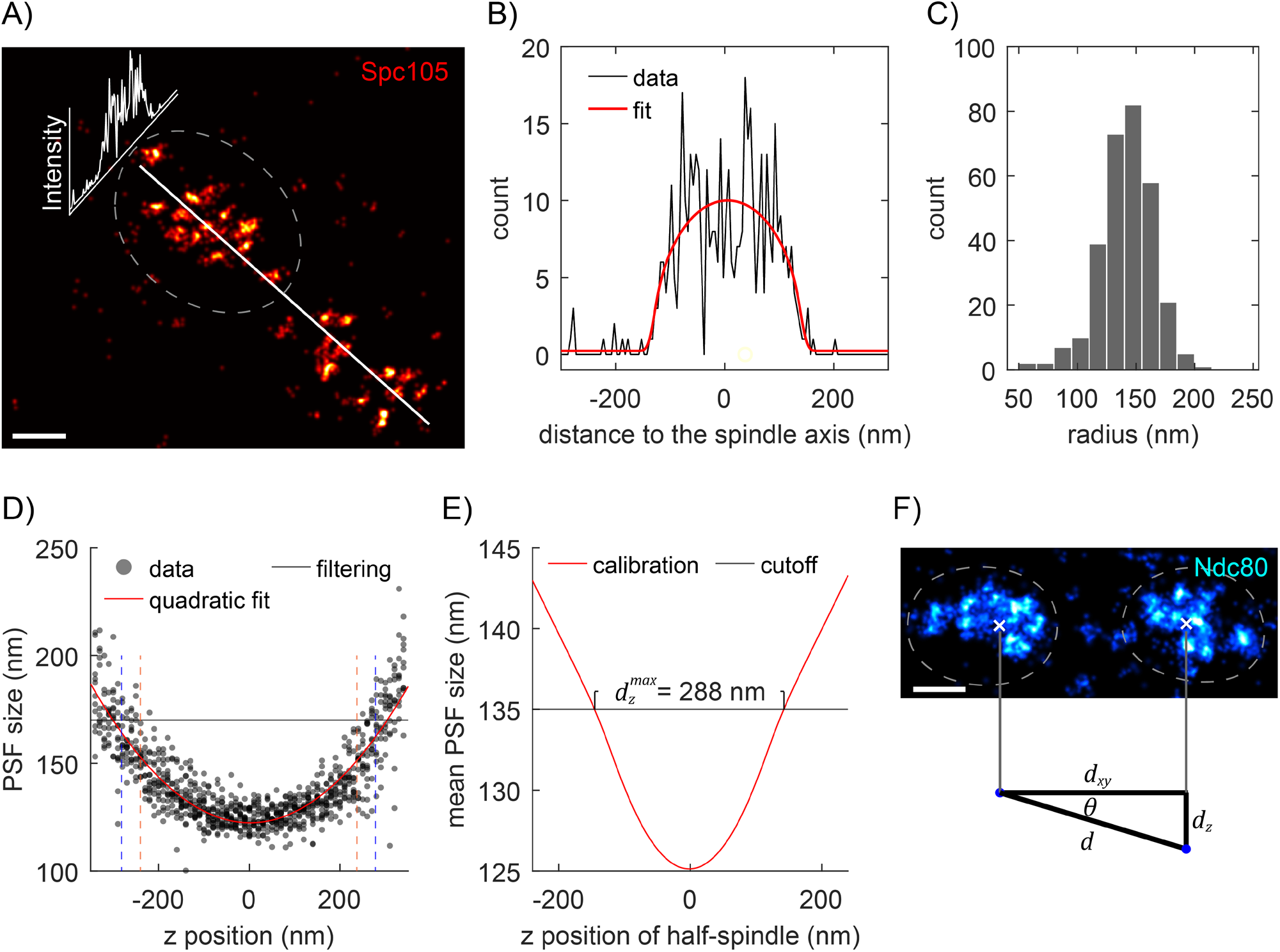
The basis for defining the values for filtering and quality control. ***A-C**. **Quantifying the width of kinetochore clusters.** As shown with the example kinetochore cluster (A), its profile perpendicular to the axis of spindle (B) was fitted with a cylindrical model (red) to quantify the radius. **C. The radius of analyzed kinetochore clusters**. The mean radius was quantified as 142.0 ± 23.7 (standard deviation) nm, which corresponds to the width (diameter) of 284 nm. Sample size: 301 kinetochore clusters. **D**. **The calibration curve (red) relating z positions to PSF size based on bead data (dots)**. For filtering out out-of-focus localizations, the maximum PSF size of 170 nm is defined, which corresponds to an axial range from −300 to 300 nm. The z ranges bounded by the vertical dashed lines with the same colors [mean PSF size cutoff: 130 nm (orange), 135 nm (blue)] are where kinetochore proteins can be found, given the corresponding mean PSF size cutoffs of kinetochore clusters, taking the quantified width in (C) into account. Both cutoffs ensure that no analyzed kinetochore protein exceeds the imaging depth determined by the PSF size filtering. **E. The calibration curve relating the z position of a kinetochore cluster to its mean PSF size**, based on the bead calibration in (D). The maximal axial distance between kinetochore clusters in the same pairs* 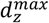 *is estimated to be 288 nm, given that the maximal allowed mean PSF size is 135 nm. **F. The relation between the lateral distance d**_xy_**, the axial distance d**_z_, **and the estimated distance between kinetochore clusters in the same pairs d in 3D**. Based on the dataset (Ndc80) with the largest sample size, the mean lateral distance between kinetochore clusters in the same pairs* 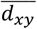 *is measured as 777 nm. These correspond to the maximum tilt angle θ^max^* = 20.3°, *and the maximum tilt-introduced error of the distance between the kinetochore clusters ϵ^max^* = 6.3%, *and the mean error* 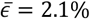. *See Methods for the calculations. Sample size: 50 kinetochore clusters. Scale bars: 200 nm*.

**Figure S2.**
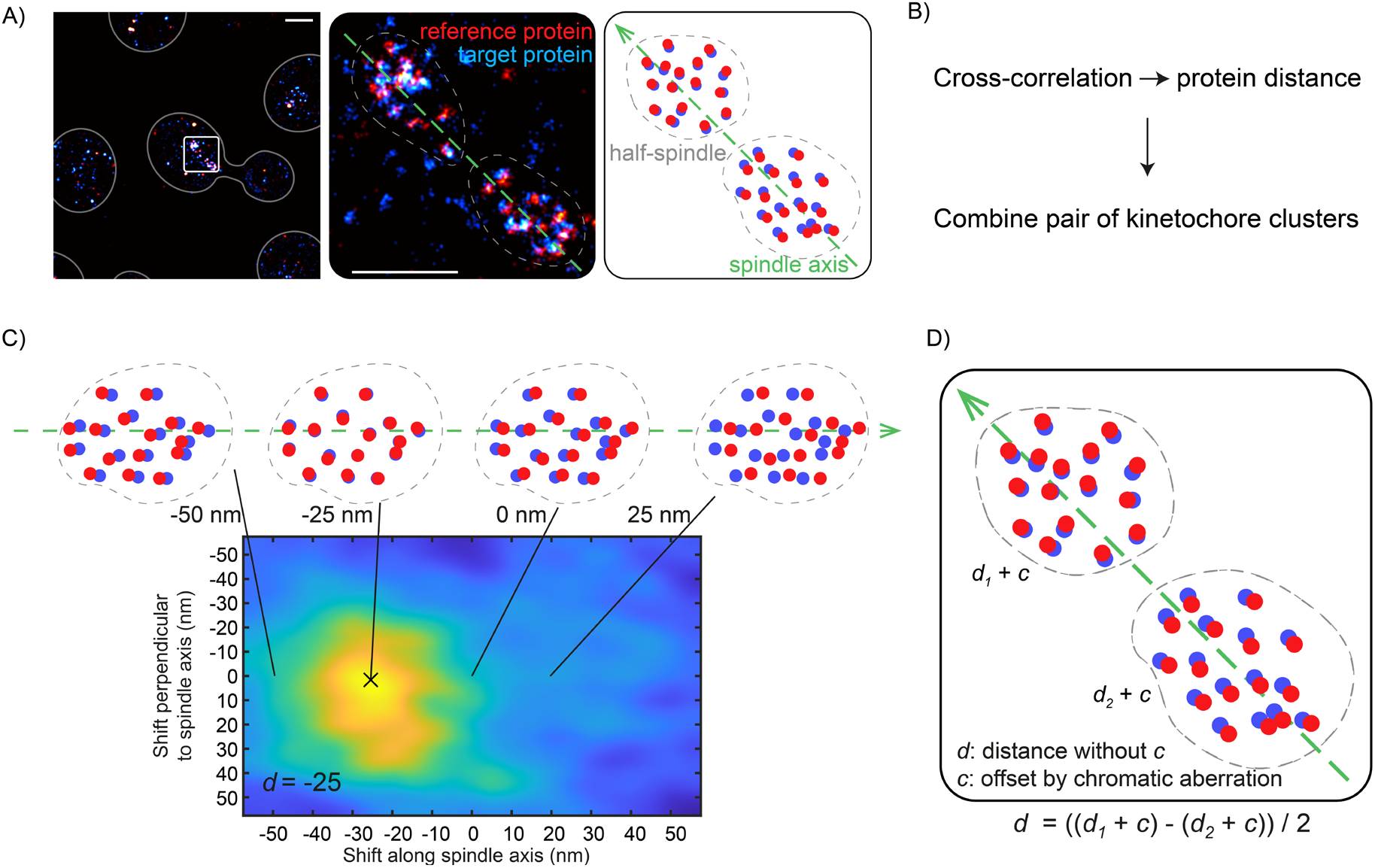
Workflow of quantifying the distances between kinetochore proteins. **A.** Metaphase spindles (white box) with both half spindles close to the focus are manually segmented (dashed contour). The spindle axis for each spindle is manually annotated (green dashed line). A schematic (right panel) is provided for clarity. **B.** The overview of the workflow. **C.** The distance between the target and reference proteins is quantified using the cross-correlation analysis. This analysis is applied to each kinetochore cluster and yields a correlation map showing the similarity between the two channels at certain lateral and axial shifts of the reference channel. The shift along spindle axis at the maximum is quantified as the distance d. **D.** To eliminate the potential offset c caused by the chromatic aberration, the average distances d of both paired kinetochore clusters, having the distances d1 and d2 respectively, is then calculated per spindle.

**Figure S3.**
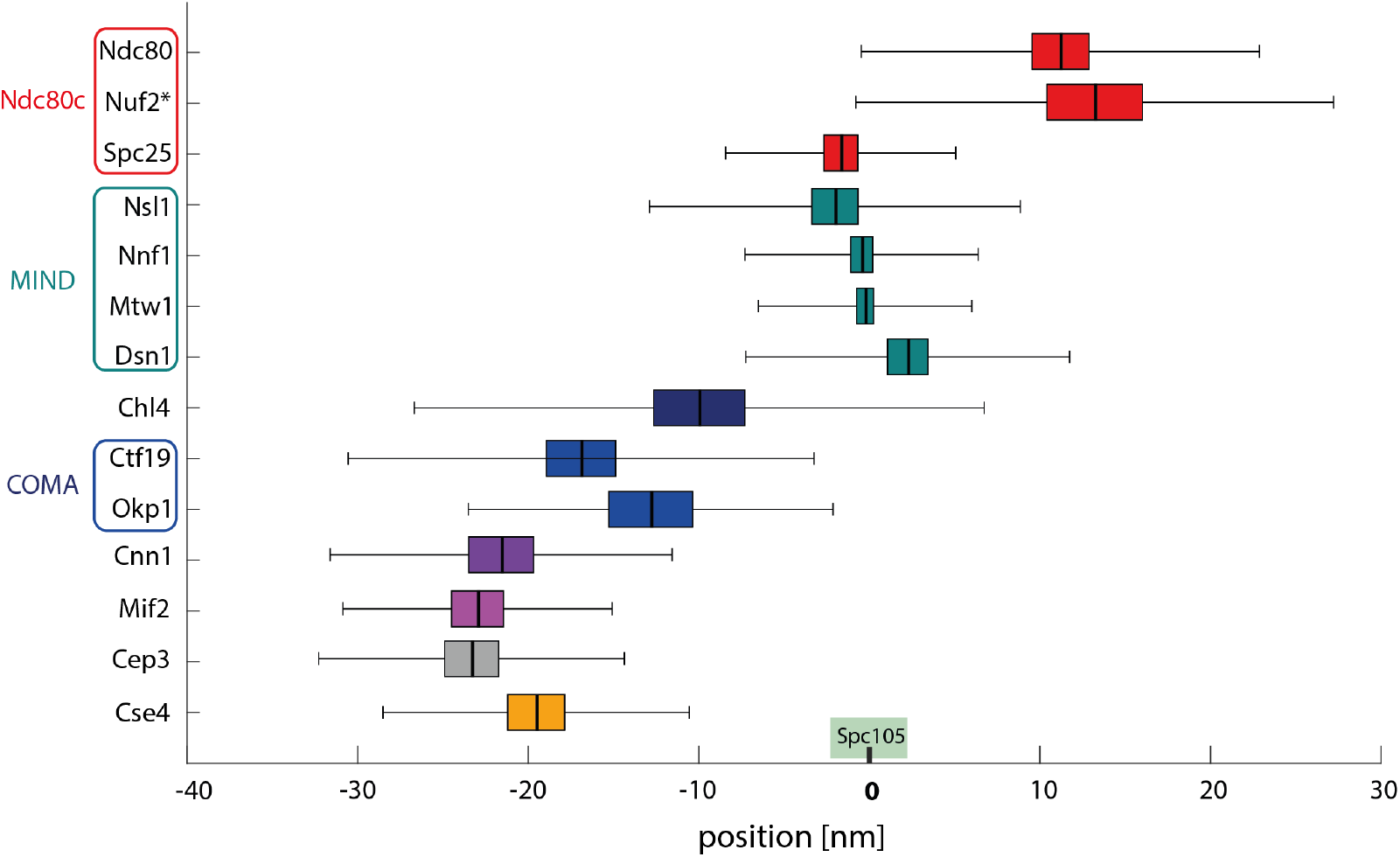
An independent analysis of intra-kinetochore distances based on manually picked single kinetochores. The mean distance is plotted with standard error of the mean - SEM (as colored box) and standard deviation - SD (whiskers). Nuf2* - the position of Nuf2 was estimated based on Nuf2-Ndc80 distance measurements.

**Figure S4.**
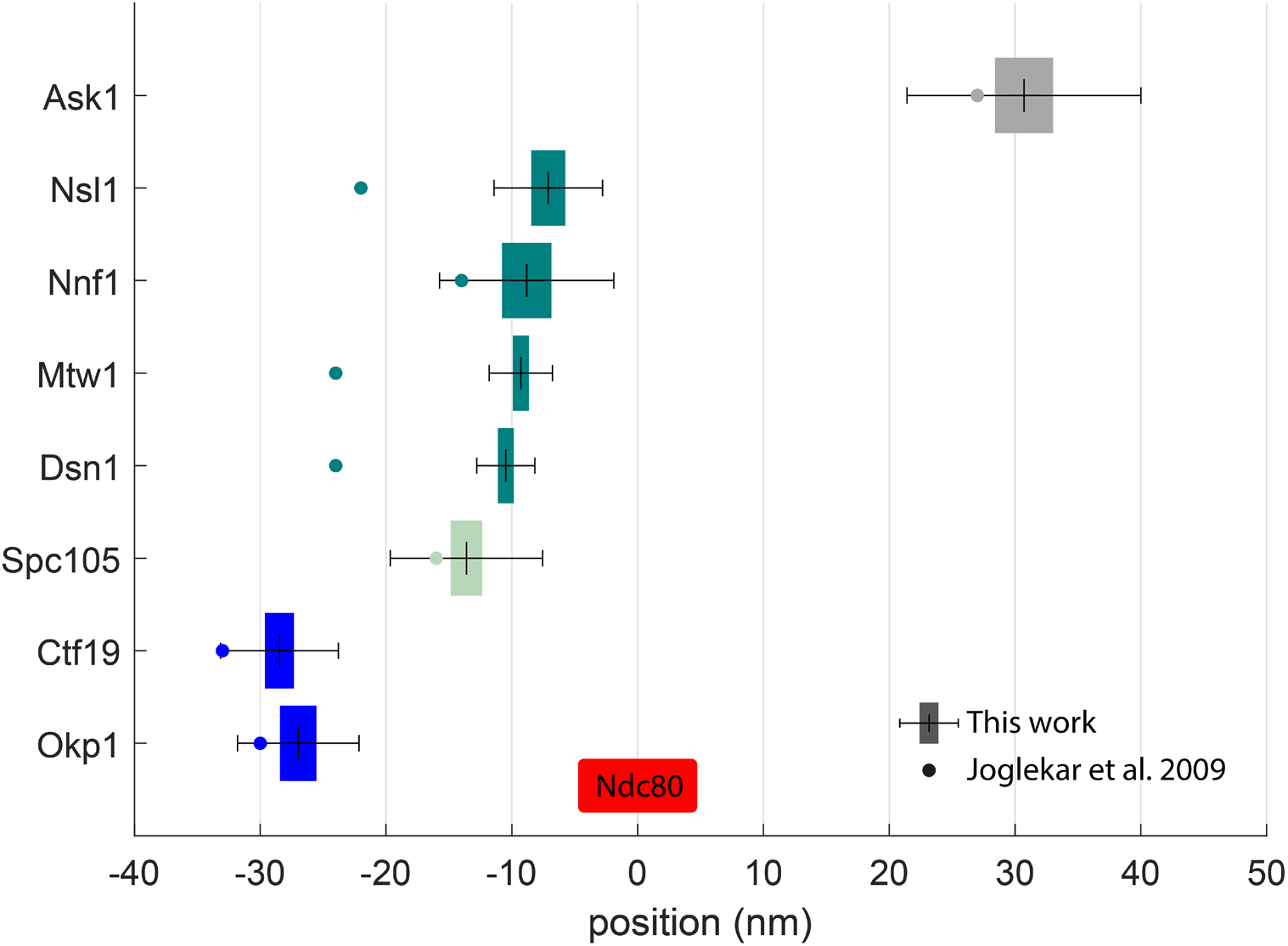
Comparison of the available distance measurements to Joglekar et al. 2009. The mean distance is plotted with standard error of the mean - SEM (as colored box) and standard deviation - SD (whiskers). The corresponding mean values reported by Joglekar et al. 2009 are shown as dots.

**Figure S5.**
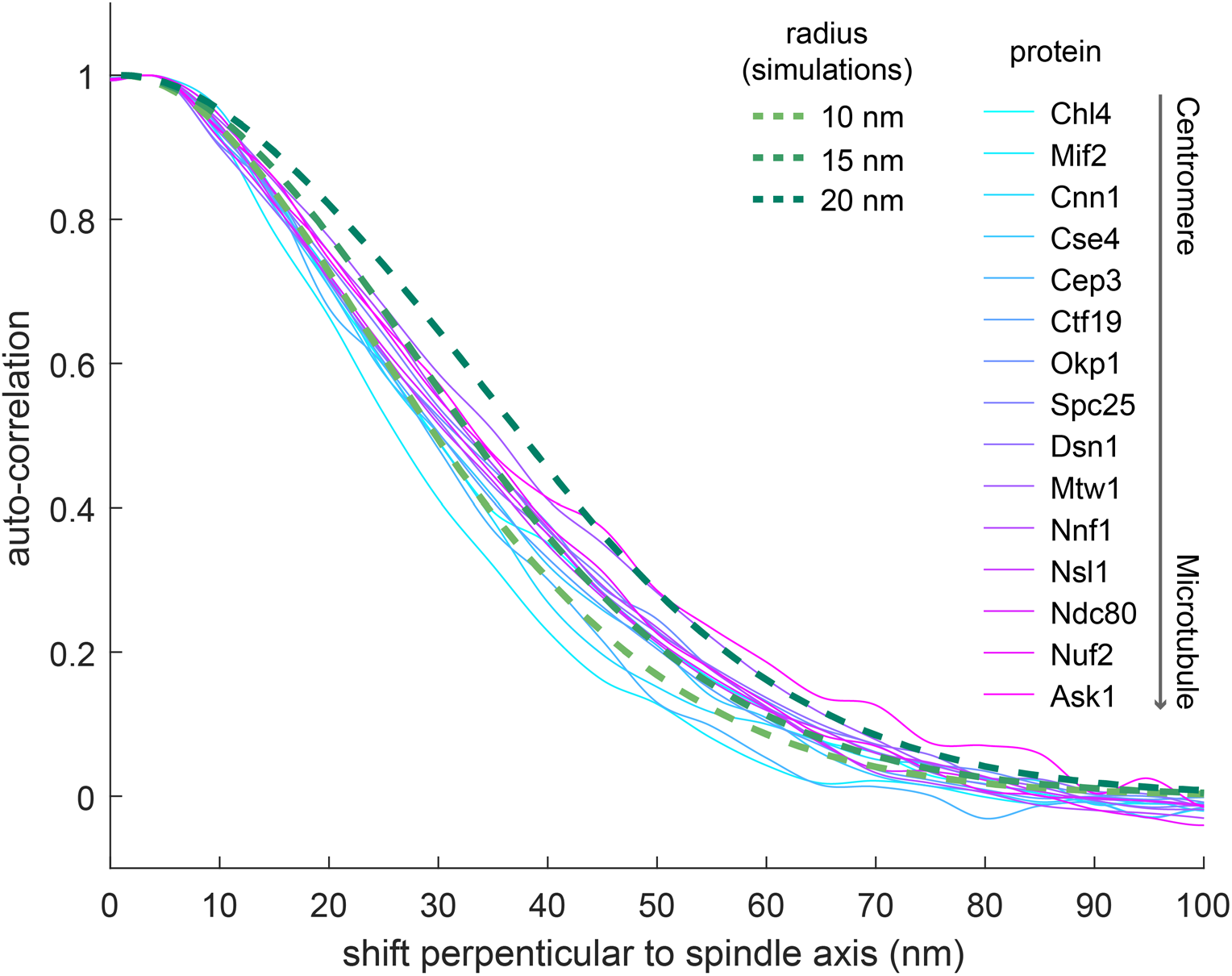
Auto-correlation perpendicular to the spindle axis. Solid curves are average auto-correlation profiles of kinetochore proteins. Dashed lines are auto-correlation profiles of simulated ring distributions with corresponding radii, considering the overall distribution of the experimental localization precision.

**Figure S6.**
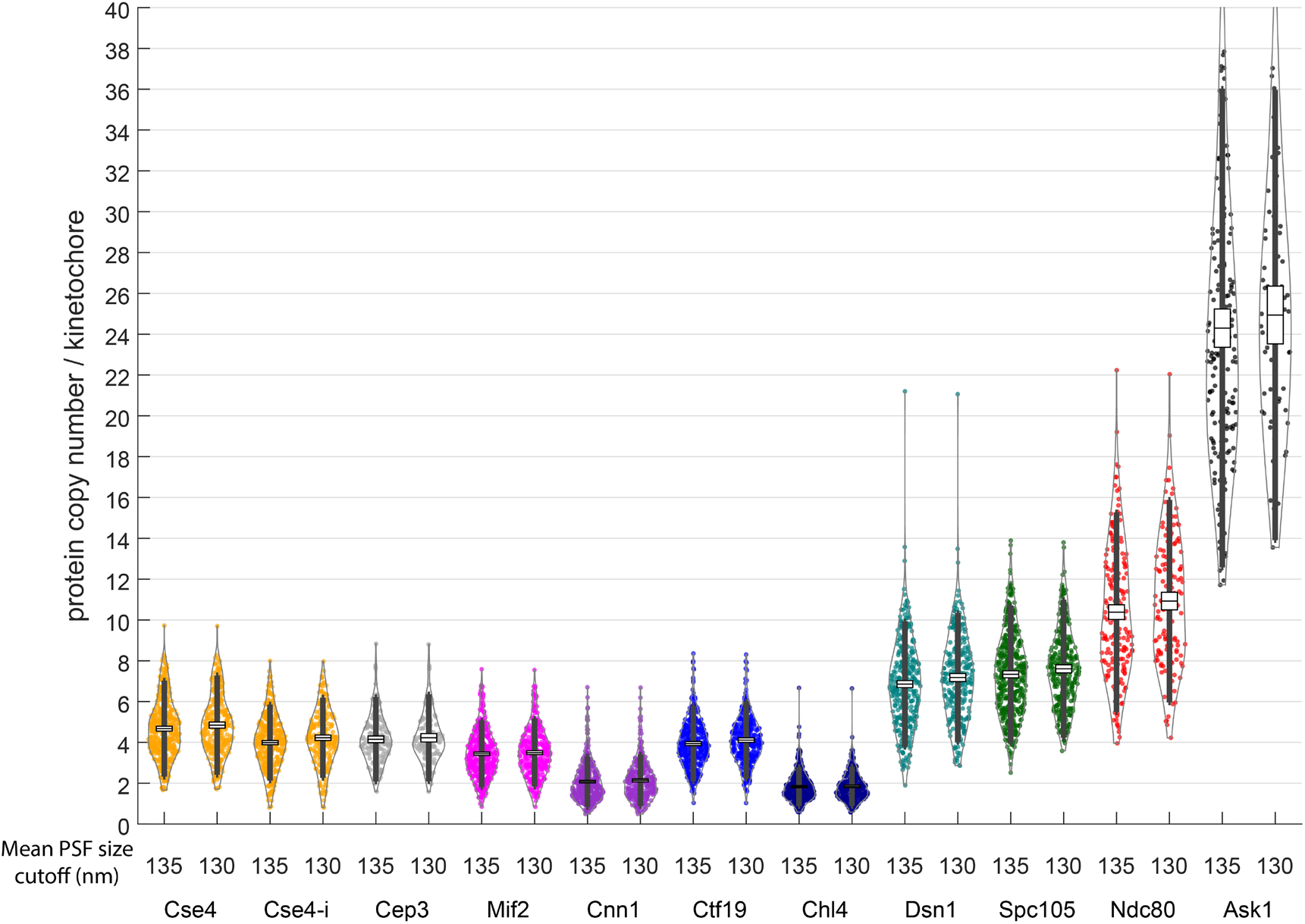
Protein copy numbers per kinetochore measured with different mean PSF size cutoffs of kinetochore clusters. (135 and 130 nm) to investigate the robustness of the molecular counting. The mean protein copy numbers calculated based on both cutoffs are almost identical, showing that the analysis is robust. Each data point corresponds to one kinetochore cluster. Boxes denote average copy numbers and standard error of means, and whiskers denote standard deviations.

**Figure S7.**
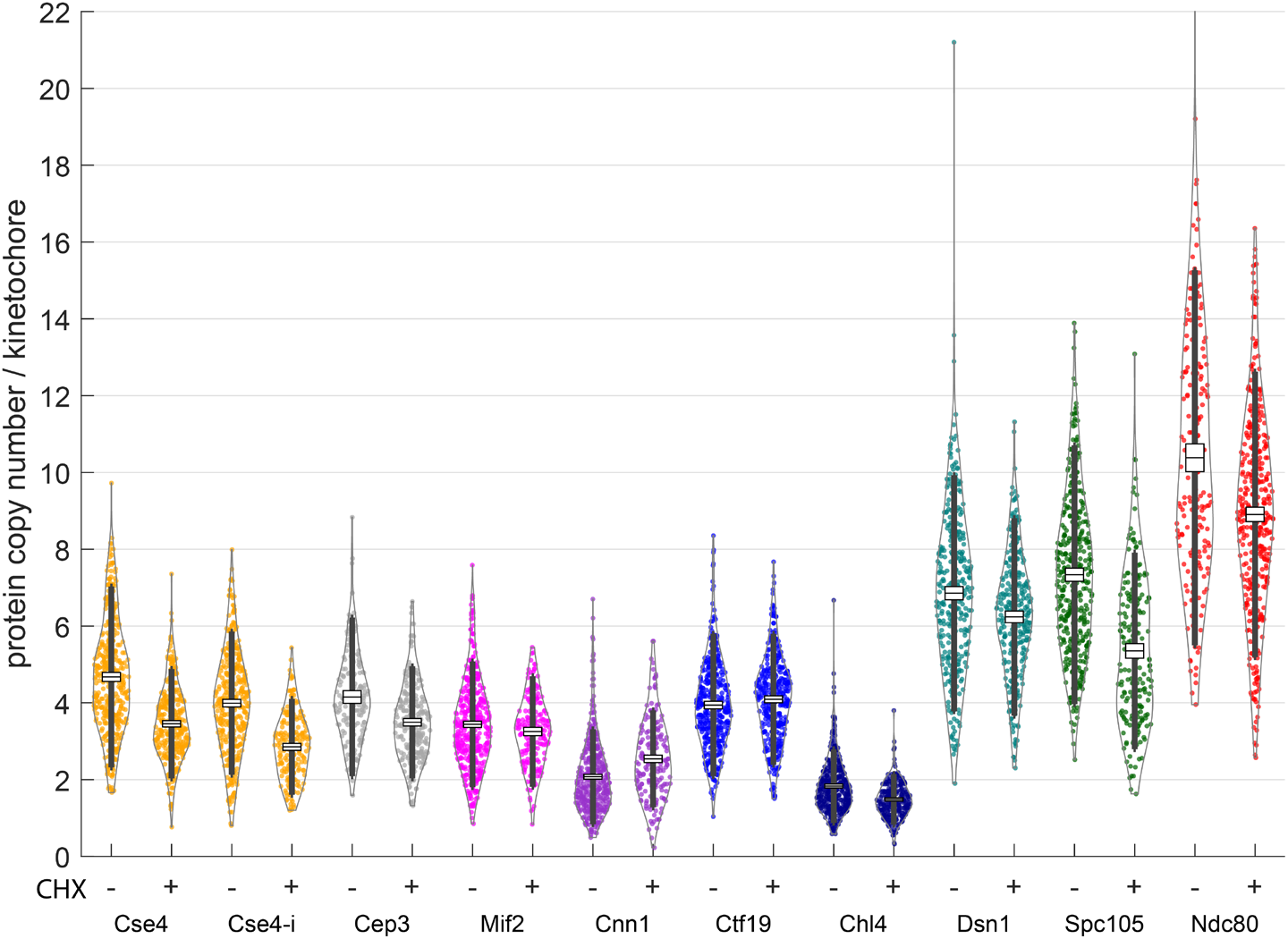
Protein copy numbers per kinetochore measured with and without cycloheximide (CHX) treatment. (250 ug/ml, 60 min), respectively, to investigate the effect of protein maturation. Each data point corresponds to one kinetochore cluster. Boxes denote average copy numbers and standard error of means, and whiskers denote standard deviations.

## Supplementary Tables

**Table S1.** Comparison of the available distance measurements from this article and Joglekar et al. 2009. Due to differences in reference points all distances were unified to the distance from Ndc80 C-terminus for clarity.

| Protein pair (with Ndc80) | Distances (nm) |  |
| --- | --- | --- |
|  | This work | Joglekar et al. 2009 |
| Ask1 | 30.7 | 27 |
| Spc105 | -13.6 | -16 |
| Nsl1 | -7.1 | -22 |
| Nnf1 | -8.8 | -14 |
| Mtw1 | -9.3 | -24 |
| Dsn1 | -10.5 | -24 |
| Ctf19 | -28.5 | -33 |
| Okp1 | -27.0 | -30 |

**Table S2.**
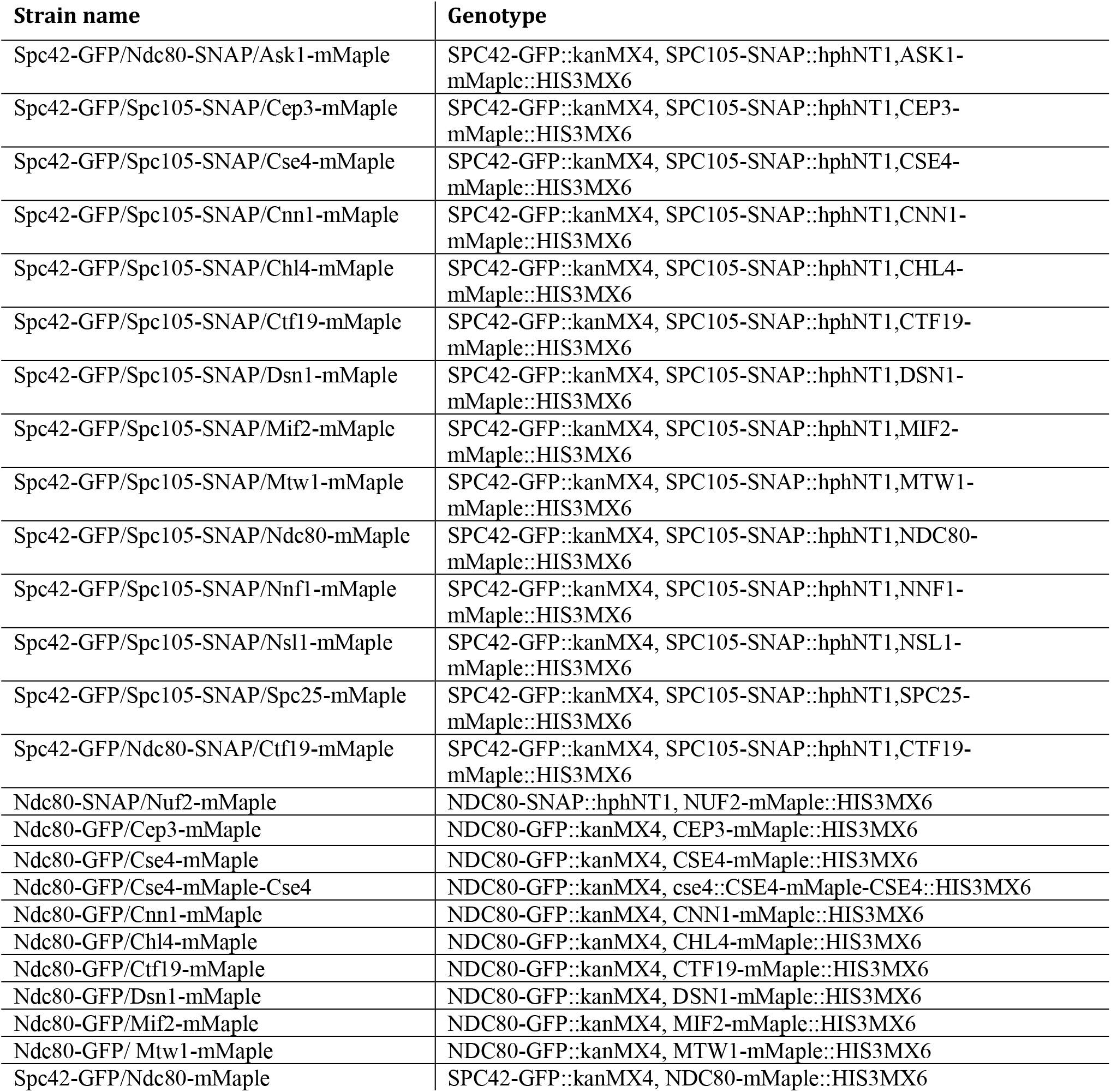
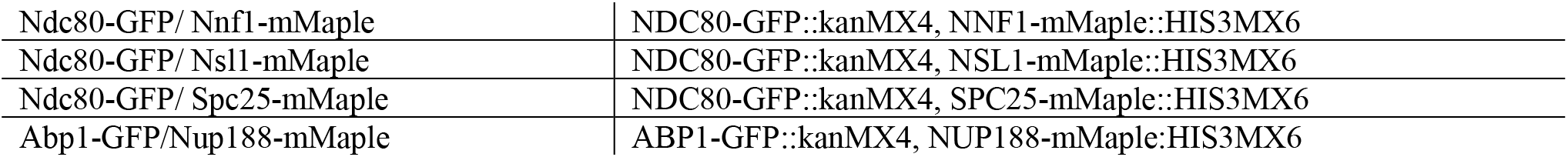
Table S2 The table represents the yeast strains created and used in this study. All are based on the MKY100 strain (S288c derivative; Kaksonen Lab) with the following genetic background: MATa, ura3-52, his3Λ200, leu3-52, lys2-801.

**Table S3.**
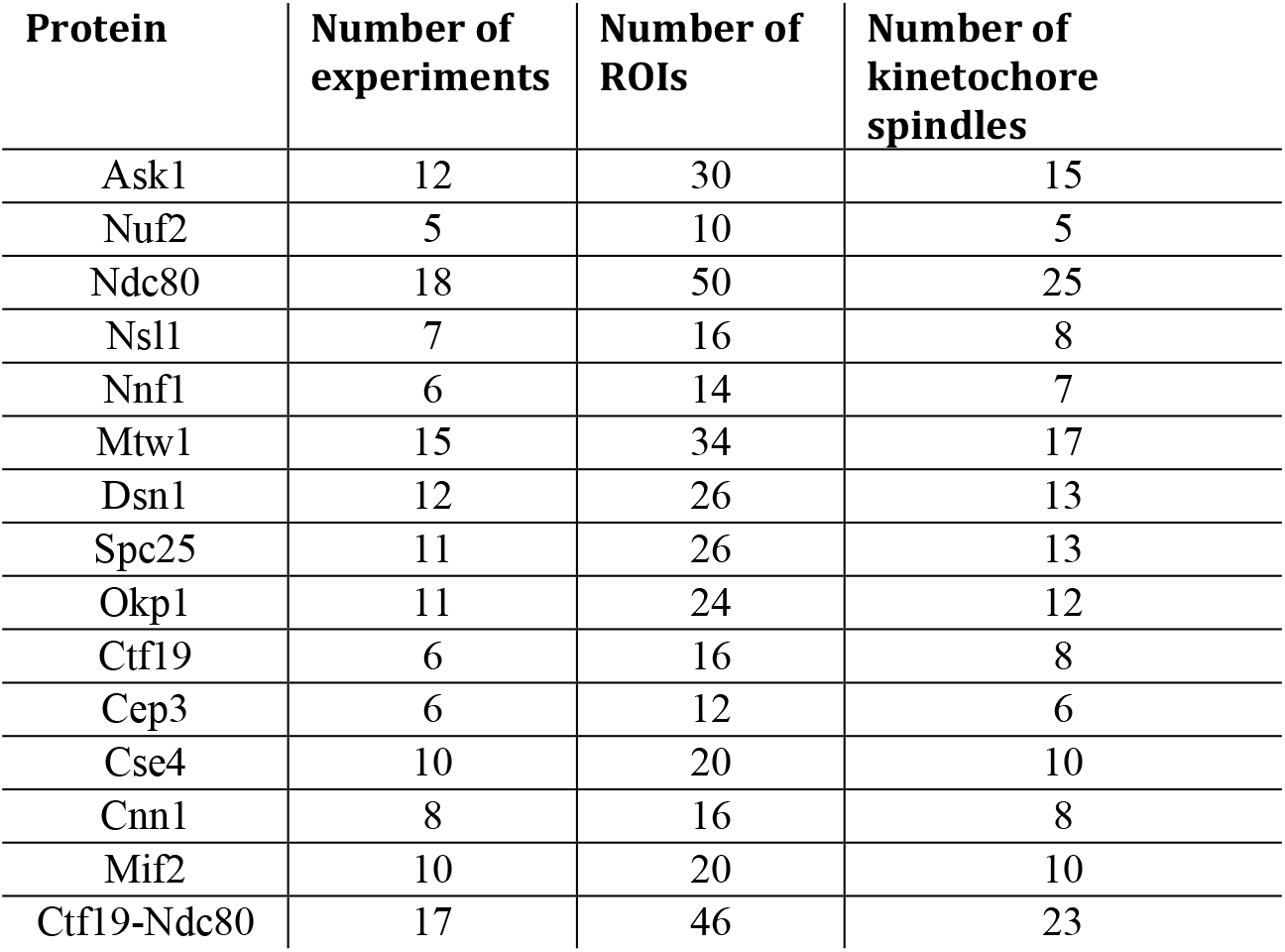
Additional information about the dual-color SMLM experiments. For each protein of interest, the number of performed experiments, ROIs and kinetochore spindles are depicted.

| Protein | Number of experiments | Number of ROIs | Number of kinetochore spindles |
| --- | --- | --- | --- |
| Ask1 | 12 | 30 | 15 |
| Nuf2 | 5 | 10 | 5 |
| Ndc80 | 18 | 50 | 25 |
| Nsl1 | 7 | 16 | 8 |
| Nnf1 | 6 | 14 | 7 |
| Mtw1 | 15 | 34 | 17 |
| Dsn1 | 12 | 26 | 13 |
| Spc25 | 11 | 26 | 13 |
| Okp1 | 11 | 24 | 12 |
| Ctf19 | 6 | 16 | 8 |
| Cep3 | 6 | 12 | 6 |
| Cse4 | 10 | 20 | 10 |
| Cnn1 | 8 | 16 | 8 |
| Mif2 | 10 | 20 | 10 |
| Ctf19-Ndc80 | 17 | 46 | 23 |

**Table S4.** Calibration factors for protein counting. The factor is the ratio between number of localizations and the copy number of Nup188 (16 copies) per NPC.

| Protein | Replicate | Calibration factor |
| --- | --- | --- |
| Ask1 | 1 | 0.89 |
|  | 2 | 1.02 |
| Ndc80 | 1 | 1.22 |
|  | 2 | 1.56 |
| Spc105 | 1 | 1.26 |
|  | 2 | 1.29 |
| Dsn1 | 1 | 1.15 |
|  | 2 | 1.51 |
| Chl4 | 1 | 1.42 |
|  | 2 | 1.17 |
| Ctf19 | 1 | 1.20 |
|  | 2 | 1.50 |
| Cnn1 | 1 | 1.14 |
|  | 2 | 1.34 |
| Mif2 | 1 | 1.15 |
|  | 2 | 1.25 |
| Cep3 | 1 | 1.46 |
|  | 2 | 1.22 |
| Cse4-i | 1 | 1.38 |
|  | 2 | 1.24 |
| Cse4 | 1 | 1.20 |
|  | 2 | 1.23 |

## References

Abbe, E., 1873. Beiträge zur Theorie des Mikroskops und der mikroskopischen Wahrnehmung. Arch. Für Mikrosk. Anat. 9, 413–418.

Ali-Ahmad, A., Bilokapić, S., Schäfer, I.B., Halić, M., Sekulić, N., 2019. CENP-C unwraps the human CENP-A nucleosome through the H2A C-terminal tail. EMBO Rep. 20, e48913. https://doi.org/10.15252/embr.201948913

Aravamudhan, P., Felzer-Kim, I., Gurunathan, K., Joglekar, A.P., 2014. Assembling the Protein Architecture of the Budding Yeast Kinetochore-Microtubule Attachment using FRET. Curr. Biol. 24, 1437–1446. https://doi.org/10.1016/j.cub.2014.05.014

Aravamudhan, P., Goldfarb, A.A., Joglekar, A.P., 2015. The kinetochore encodes a mechanical switch to disrupt spindle assembly checkpoint signalling. Nat. Cell Biol. 17, 868–879. https://doi.org/10.1038/ncb3179

Asbury, C.L., 2017. Anaphase A: Disassembling Microtubules Move Chromosomes toward Spindle Poles. Biology 6, 15. https://doi.org/10.3390/biology6010015

Betzig, E., Patterson, G.H., Sougrat, R., Lindwasser, O.W., Olenych, S., Bonifacino, J.S., Davidson, M.W., Lippincott-Schwartz, J., Hess, H.F., 2006. Imaging Intracellular Fluorescent Proteins at Nanometer Resolution. Science 313, 1642–1645. https://doi.org/10/dd8dkd

Black, B.E., Cleveland, D.W., 2011. Epigenetic Centromere Propagation and the Nature of CENP-A Nucleosomes. Cell 144, 471–479. https://doi.org/10.1016/j.cell.2011.02.002

Bui, M., Dimitriadis, E.K., Hoischen, C., An, E., Quénet, D., Giebe, S., Nita-Lazar, A., Diekmann, S., Dalal, Y., 2012. Cell-Cycle-Dependent Structural Transitions in the Human CENP-A Nucleosome In Vivo. Cell 150, 317–326. https://doi.org/10.1016/j.cell.2012.05.035

Bystricky, K., Laroche, T., van Houwe, G., Blaszczyk, M., Gasser, S.M., 2005. Chromosome looping in yeast telomere pairing and coordinated movement reflect anchoring efficiency and territorial organization. J. Cell Biol. 168, 375–387. https://doi.org/10.1083/jcb.200409091

Camahort, R., Shivaraju, M., Mattingly, M., Li, B., Nakanishi, S., Zhu, D., Shilatifard, A., Workman, J.L., Gerton, J.L., 2009. Cse4 Is Part of an Octameric Nucleosome in Budding Yeast. Mol. Cell 35, 794–805. https://doi.org/10.1016/j.molcel.2009.07.022

Carroll, C.W., Milks, K.J., Straight, A.F., 2010. Dual recognition of CENP-A nucleosomes is required for centromere assembly. J. Cell Biol. 189, 1143–1155. https://doi.org/10.1083/jcb.201001013

Chaly, N., Brown, D.L., 1988. The prometaphase configuration and chromosome order in early mitosis. J. Cell Sci. 91 (Pt 3), 325–335. https://doi.org/10.1242/jcs.91.3.325

Cheeseman, I.M., Chappie, J.S., Wilson-Kubalek, E.M., Desai, A., 2006. The Conserved KMN Network Constitutes the Core Microtubule-Binding Site of the Kinetochore. Cell 127, 983–997. https://doi.org/10.1016/j.cell.2006.09.039

Ciferri, C., Pasqualato, S., Screpanti, E., Varetti, G., Santaguida, S., Dos Reis, G., Maiolica, A., Polka, J., De Luca, J.G., De Wulf, P., Salek, M., Rappsilber, J., Moores, C.A., Salmon, E.D., Musacchio, A., 2008. Implications for Kinetochore-Microtubule Attachment from the Structure of an Engineered Ndc80 Complex. Cell 133, 427–439. https://doi.org/10.1016/j.cell.2008.03.020

Clarke, L., Carbon, J., 1980. Isolation of a yeast centromere and construction of functional small circular chromosomes. Nature 287, 504–509. https://doi.org/10.1038/287504a0

Cohen, R.L., Espelin, C.W., De Wulf, P., Sorger, P.K., Harrison, S.C., Simons, K.T., 2008. Structural and Functional Dissection of Mif2p, a Conserved DNA-binding Kinetochore Protein. Mol. Biol. Cell 19, 4480–4491. https://doi.org/10.1091/mbc.e08-03-0297

Conti, D., Hart, M., Tamura, N., Shrestha, R., Islam, A., Draviam, V.M., 2017. How are Dynamic Microtubules Stably Tethered to Human Chromosomes? Cytoskelet. - Struct. Dyn. Funct. Dis. https://doi.org/10.5772/intechopen.68321

Dalal, Y., Furuyama, T., Vermaak, D., Henikoff, S., 2007. Structure, dynamics, and evolution of centromeric nucleosomes. Proc. Natl. Acad. Sci. 104, 15974. https://doi.org/10.1073/pnas.0707648104

Dani, A., Huang, B., Bergan, J., Dulac, C., Zhuang, X., 2010. Superresolution Imaging of Chemical Synapses in the Brain. Neuron 68, 843–856. https://doi.org/10.1016/j.neuron.2010.11.021

Deschamps, J., Ries, J., 2020. EMU: reconfigurable graphical user interfaces for Micro-Manager. BMC Bioinformatics 21, 456. https://doi.org/10.1186/s12859-020-03727-8

Deschamps, J., Rowald, A., Ries, J., 2016. Efficient homogeneous illumination and optical sectioning for quantitative single-molecule localization microscopy. Opt. Express 24, 28080–28090.

Dhatchinamoorthy, K., Shivaraju, M., Lange, J.J., Rubinstein, B., Unruh, J.R., Slaughter, B.D., Gerton, J.L., 2017. Structural plasticity of the living kinetochore. J. Cell Biol. 216, 3551–3570. https://doi.org/10.1083/jcb.201703152

Dimitrova, Y.N., Jenni, S., Valverde, R., Khin, Y., Harrison, S.C., 2016. Structure of the MIND Complex Defines a Regulatory Focus for Yeast Kinetochore Assembly. Cell 167, 1014–1027. https://doi.org/10.1016/j.cell.2016.10.011

Drinnenberg, I.A., Henikoff, S., Malik, H.S., 2016. Evolutionary Turnover of Kinetochore Proteins: A Ship of Theseus? Trends Cell Biol. 26, 498–510. https://doi.org/10.1016/j.tcb.2016.01.005

Edelstein, A.D., Tsuchida, M.A., Amodaj, N., Pinkard, H., Vale, R.D., Stuurman, N., 2014. Advanced methods of microscope control using μManager software. J. Biol. Methods 1, e10. https://doi.org/10.14440/jbm.2014.36

Fischböck-Halwachs, J., Singh, S., Potocnjak, M., Hagemann, G., Solis-Mezarino, V., Woike, S., Ghodgaonkar-Steger, M., Weissmann, F., Gallego, L.D., Rojas, J., Andreani, J., Köhler, A., Herzog, F., 2019. The COMA complex interacts with Cse4 and positions Sli15/Ipl1 at the budding yeast inner kinetochore. eLife 8, e42879. https://doi.org/10.7554/eLife.42879

Furuyama, S., Biggins, S., 2007. Centromere identity is specified by a single centromeric nucleosome in budding yeast. Proc. Natl. Acad. Sci. 104, 14706–14711. https://doi.org/10.1073/pnas.0706985104

Gonen, S., Akiyoshi, B., Iadanza, M.G., Shi, D., Duggan, N., Biggins, S., Gonen, T., 2012. The structure of purified kinetochores reveals multiple microtubule-attachment sites. Nat. Struct. Mol. Biol. 19, 925–929. https://doi.org/10.1038/nsmb.2358

Guse, A., Carroll, C.W., Moree, B., Fuller, C.J., Straight, A.F., 2011. In vitro centromere and kinetochore assembly on defined chromatin templates. Nature 477, 354–358. https://doi.org/10.1038/nature10379

Haase, J., Mishra, P.K., Stephens, A., Haggerty, R., Quammen, C., Taylor, R.M., Yeh, E., Basrai, M.A., Bloom, K., 2013. A 3D Map of the Yeast Kinetochore Reveals the Presence of Core and Accessory Centromere-Specific Histone. Curr. Biol. 23, 1939–1944. https://doi.org/10.1016/j.cub.2013.07.083

Hamilton, G., Dimitrova, Y., Davis, T.N., 2019. Seeing is believing: our evolving view of kinetochore structure, composition, and assembly. Curr. Opin. Cell Biol. 60, 44–52. https://doi.org/10.1016/j.ceb.2019.03.016

Hess, S.T., Girirajan, T.P.K., Mason, M.D., 2006. Ultra-High Resolution Imaging by Fluorescence Photoactivation Localization Microscopy. Biophys. J. 91, 4258–4272. https://doi.org/10/cr8s93

Hinshaw, S.M., Harrison, S.C., 2019. The structure of the Ctf19c/CCAN from budding yeast. eLife 8, e44239. https://doi.org/10.7554/eLife.44239

Hooff, J.J., Tromer, E., Wijk, L.M., Snel, B., Kops, G.J., 2017. Evolutionary dynamics of the kinetochore network in eukaryotes as revealed by comparative genomics. EMBO Rep. 18, 1559–1571. https://doi.org/10.15252/embr.201744102

Hornung, P., Maier, M., Alushin, G.M., Lander, G.C., Nogales, E., Westermann, S., 2011. Molecular Architecture and Connectivity of the Budding Yeast Mtw1 Kinetochore Complex. J. Mol. Biol. 405, 548–559. https://doi.org/10.1016/j.jmb.2010.11.012

Hornung, P., Troc, P., Malvezzi, F., Maier, M., Demianova, Z., Zimniak, T., Litos, G., Lampert, F., Schleiffer, A., Brunner, M., Mechtler, K., Herzog, F., Marlovits, T.C., Westermann, S., 2014. A cooperative mechanism drives budding yeast kinetochore assembly downstream of CENP-A. J. Cell Biol. 206, 509–524. https://doi.org/10.1083/jcb.201403081

Huis in ’t Veld, P.J., Jeganathan, S., Petrovic, A., Singh, P., John, J., Krenn, V., Weissmann, F., Bange, T., Musacchio, A., 2016. Molecular basis of outer kinetochore assembly on CENP-T. eLife 5, e21007. https://doi.org/10.7554/eLife.21007

Janke, C., Magiera, M.M., Rathfelder, N., Taxis, C., Reber, S., Maekawa, H., Moreno-Borchart, A., Doenges, G., Schwob, E., Schiebel, E., Knop, M., 2004. A versatile toolbox for PCR-based tagging of yeast genes: new fluorescent proteins, more markers and promoter substitution cassettes. Yeast 21, 947–962. https://doi.org/10.1002/yea.1142

Jenni, S., Dimitrova, Y.N., Valverde, R., Hinshaw, S.M., Harrison, S.C., 2017. Molecular Structures of Yeast Kinetochore Subcomplexes and Their Roles in Chromosome Segregation. Cold Spring Harb. Symp. Quant. Biol. 82, 83–89. https://doi.org/10.1101/sqb.2017.82.033738

Jenni, S., Harrison, S.C., 2018. Structure of the DASH/Dam1 complex shows its role at the yeast kinetochore-microtubule interface. Science 360, 552–558. https://doi.org/10.1126/science.aar6436

Jin, Q.W., Fuchs, J., Loidl, J., 2000. Centromere clustering is a major determinant of yeast interphase nuclear organization. J. Cell Sci. 113, 1903–1912.

Joglekar, A.P., Bloom, K., Salmon, E.D., 2009. In Vivo Protein Architecture of the Eukaryotic Kinetochore with Nanometer Scale Accuracy. Curr. Biol. 19, 694–699. https://doi.org/10.1016/j.cub.2009.02.056

Joglekar, A.P., Bloom, K.S., Salmon, E., 2010. Mechanisms of force generation by end-on kinetochore-microtubule attachments. Curr. Opin. Cell Biol. 22, 57–67. https://doi.org/10.1016/j.ceb.2009.12.010

Joglekar, A.P., Bouck, D.C., Molk, J.N., Bloom, K.S., Salmon, E.D., 2006. Molecular architecture of a kinetochore– microtubule attachment site. Nat. Cell Biol. 8, 581–585. https://doi.org/10.1038/ncb1414

Joglekar, A.P., Salmon, E.D., Bloom, K.S., 2008. Counting Kinetochore Protein Numbers in Budding Yeast Using Genetically Encoded Fluorescent Proteins. Methods Cell Biol. 85, 127–151. https://doi.org/10.1016/S0091-679X(08)85007-8

Keith, K.C., Fitzgerald-Hayes, M., 2000. CSE4 Genetically Interacts With the *Saccharomyces cerevisiae* Centromere DNA Elements CDE I and CDE II but Not CDE III: Implications for the Path of the Centromere DNA Around a Cse4p Variant Nucleosome. Genetics 156, 973–981. https://doi.org/10.1093/genetics/156.3.973

Kim, J. ook, Zelter, A., Umbreit, N.T., Bollozos, A., Riffle, M., Johnson, R., MacCoss, M.J., Asbury, C.L., Davis, T.N., 2017. The Ndc80 complex bridges two Dam1 complex rings. eLife 6, e21069. https://doi.org/10.7554/eLife.21069

Kim, S.J., Fernandez-Martinez, J., Nudelman, I., Shi, Y., Zhang, W., Raveh, B., Herricks, T., Slaughter, B.D., Hogan, J.A., Upla, P., Chemmama, I.E., Pellarin, R., Echeverria, I., Shivaraju, M., Chaudhury, A.S., Wang, J., Williams, R., Unruh, J.R., Greenberg, C.H., Jacobs, E.Y., Yu, Z., de la Cruz, M.J., Mironska, R., Stokes, D.L., Aitchison, J.D., Jarrold, M.F., Gerton, J.L., Ludtke, S.J., Akey, C.W., Chait, B.T., Sali, A., Rout, M.P., 2018. Integrative structure and functional anatomy of a nuclear pore complex. Nature 555, 475–482. https://doi.org/10.1038/nature26003

Krassovsky, K., Henikoff, J.G., Henikoff, S., 2012. Tripartite organization of centromeric chromatin in budding yeast. Proc. Natl. Acad. Sci. 109, 243–248. https://doi.org/10.1073/pnas.1118898109

Kudalkar, E.M., Scarborough, E.A., Umbreit, N.T., Zelter, A., Gestaut, D.R., Riffle, M., Johnson, R.S., MacCoss, M.J., Asbury, C.L., Davis, T.N., 2015. Regulation of outer kinetochore Ndc80 complex-based microtubule attachments by the central kinetochore Mis12/MIND complex. Proc. Natl. Acad. Sci. 112, E5583–E5589. https://doi.org/10.1073/pnas.1513882112

Lando, D., Endesfelder, U., Berger, H., Subramanian, L., Dunne, P.D., McColl, J., Klenerman, D., Carr, A.M., Sauer, M., Allshire, R.C., Heilemann, M., Laue, E.D., 2012. Quantitative single-molecule microscopy reveals that CENP-A ^Cnp1^ deposition occurs during G2 in fission yeast. Open Biol. 2, 120078. https://doi.org/10.1098/rsob.120078

Lang, J., Barber, A., Biggins, S., 2018. An assay for de novo kinetochore assembly reveals a key role for the CENP-T pathway in budding yeast. eLife 7. https://doi.org/10.7554/eLife.37819

Lawrimore, J., Bloom, K.S., Salmon, E.D., 2011. Point centromeres contain more than a single centromere-specific Cse4 (CENP-A) nucleosome. J. Cell Biol. 195, 573–582.

Leber, V., Nans, A., Singleton, M.R., 2018. Structural basis for assembly of the CBF3 kinetochore complex. EMBO J. 37, 269–281. https://doi.org/10.15252/embj.201798134

Malvezzi, F., Litos, G., Schleiffer, A., Heuck, A., Mechtler, K., Clausen, T., Westermann, S., 2013. A structural basis for kinetochore recruitment of the Ndc80 complex via two distinct centromere receptors. EMBO J. 32, 409–423. https://doi.org/10.1038/emboj.2012.356

McEvoy, A.L., Hoi, H., Bates, M., Platonova, E., Cranfill, P.J., Baird, M.A., Davidson, M.W., Ewers, H., Liphardt, J., Campbell, R.E., 2012. mMaple: A Photoconvertible Fluorescent Protein for Use in Multiple Imaging Modalities. PLoS ONE 7, e51314. https://doi.org/10.1371/journal.pone.0051314

McIntosh, J.R., O’Toole, E., Zhudenkov, K., Morphew, M., Schwartz, C., Ataullakhanov, F.I., Grishchuk, E.L., 2013. Conserved and divergent features of kinetochores and spindle microtubule ends from five species. J. Cell Biol. 200, 459–474. https://doi.org/10.1083/jcb.201209154

McKinley, K.L., Sekulic, N., Guo, L.Y., Tsinman, T., Black, B.E., Cheeseman, I.M., 2015. The CENP-L-N Complex Forms a Critical Node in an Integrated Meshwork of Interactions at the Centromere-Kinetochore Interface. Mol. Cell 60, 886–898. https://doi.org/10.1016/j.molcel.2015.10.027

Mizuguchi, G., Xiao, H., Wisniewski, J., Smith, M.M., Wu, C., 2007. Nonhistone Scm3 and Histones CenH3-H4 Assemble the Core of Centromere-Specific Nucleosomes. Cell 129, 1153–1164. https://doi.org/10.1016/j.cell.2007.04.026

Mund, M., van der Beek, J.A., Deschamps, J., Dmitrieff, S., Hoess, P., Monster, J.L., Picco, A., Nédélec, F., Kaksonen, M., Ries, J., 2018. Systematic Nanoscale Analysis of Endocytosis Links Efficient Vesicle Formation to Patterned Actin Nucleation. Cell 174, 884–896.e17. https://doi.org/10.1016/j.cell.2018.06.032

Musacchio, A., Desai, A., 2017. A Molecular View of Kinetochore Assembly and Function. Biology 6, 5. https://doi.org/10.3390/biology6010005

Ng, C.T., Deng, L., Chen, C., Lim, H.H., Shi, J., Surana, U., Gan, L., 2018. Electron cryotomography analysis of Dam1C/DASH at the kinetochore–spindle interface in situ. J. Cell Biol. 218, 455–473. https://doi.org/10.1083/jcb.201809088

Ng, R., Carbon, J., 1987. Mutational and in vitro protein-binding studies on centromere DNA from Saccharomyces cerevisiae. Mol. Cell. Biol. 7, 4522–4534. https://doi.org/10.1128/MCB.7.12.4522

Pekgöz Altunkaya, G., Malvezzi, F., Demianova, Z., Zimniak, T., Litos, G., Weissmann, F., Mechtler, K., Herzog, F., Westermann, S., 2016. CCAN Assembly Configures Composite Binding Interfaces to Promote Cross-Linking of Ndc80 Complexes at the Kinetochore. Curr. Biol. 26, 2370–2378. https://doi.org/10.1016/j.cub.2016.07.005

Pentakota, S., Zhou, K., Smith, C., Maffini, S., Petrovic, A., Morgan, G.P., Weir, J.R., Vetter, I.R., Musacchio, A., Luger, K., 2017. Decoding the centromeric nucleosome through CENP-N. eLife 6, e33442. https://doi.org/10.7554/eLife.33442

Pesenti, M.E., Raisch, T., Conti, D., Walstein, K., Hoffmann, I., Vogt, D., Prumbaum, D., Vetter, I.R., Raunser, S., Musacchio, A., 2022. Structure of the human inner kinetochore CCAN complex and its significance for human centromere organization. Mol. Cell 82, 2113–2131. https://doi.org/10.1016/j.molcel.2022.04.027

Petrovic, A., Keller, J., Liu, Y., Overlack, K., John, J., Dimitrova, Y.N., Jenni, S., van Gerwen, S., Stege, P., Wohlgemuth, S., Rombaut, P., Herzog, F., Harrison, S.C., Vetter, I.R., Musacchio, A., 2016. Structure of the MIS12 Complex and Molecular Basis of Its Interaction with CENP-C at Human Kinetochores. Cell 167, 1028–1040. https://doi.org/10.1016/j.cell.2016.10.005

Petrovic, A., Mosalaganti, S., Keller, J., Mattiuzzo, M., Overlack, K., Krenn, V., De Antoni, A., Wohlgemuth, S., Cecatiello, V., Pasqualato, S., Raunser, S., Musacchio, A., 2014. Modular Assembly of RWD Domains on the Mis12 Complex Underlies Outer Kinetochore Organization. Mol. Cell 53, 591–605. https://doi.org/10.1016/j.molcel.2014.01.019

Przewloka, M.R., Venkei, Z., Bolanos-Garcia, V.M., Debski, J., Dadlez, M., Glover, D.M., 2011. CENP-C Is a Structural Platform for Kinetochore Assembly. Curr. Biol. 21, 399–405. https://doi.org/10.1016/j.cub.2011.02.005

Ramey, V.H., Wang, H.-W., Nakajima, Y., Wong, A., Liu, J., Drubin, D., Barnes, G., Nogales, E., 2011. The Dam1 ring binds to the E-hook of tubulin and diffuses along the microtubule. Mol. Biol. Cell 22, 457–466. https://doi.org/10.1091/mbc.E10-10-0841

Rieder, C.L., 1982. The Formation, Structure, and Composition of the Mammalian Kinetochore and Kinetochore Fiber, in: International Review of Cytology. Elsevier, pp. 1–58. https://doi.org/10.1016/S0074-7696(08)61672-1

Ries, J., 2020. SMAP: a modular super-resolution microscopy analysis platform for SMLM data. Nat. Methods 17, 870–872. https://doi.org/10.1038/s41592-020-0938-1

Rust, M.J., Bates, M., Zhuang, X., 2006. Sub-diffraction-limit imaging by stochastic optical reconstruction microscopy (STORM). Nat. Methods 3, 793–795. https://doi.org/10/fwrhq3

Santaguida, S., Amon, A., 2015. Short-and long-term effects of chromosome mis-segregation and aneuploidy. Nat. Rev. Mol. Cell Biol. 16, 473–485. https://doi.org/10.1038/nrm4025

Schmitzberger, F., Richter, M.M., Gordiyenko, Y., Robinson, C.V., Dadlez, M., Westermann, S., 2017. Molecular basis for inner kinetochore configuration through RWD domain–peptide interactions. EMBO J. 36, 3458–3482. https://doi.org/10.15252/embj.201796636

Scott, K.C., Bloom, K.S., 2014. Lessons learned from counting molecules: how to lure CENP-A into the kinetochore. Open Biol. 4, 140191. https://doi.org/10.1098/rsob.140191

Screpanti, E., De Antoni, A., Alushin, G.M., Petrovic, A., Melis, T., Nogales, E., Musacchio, A., 2011. Direct Binding of Cenp-C to the Mis12 Complex Joins the Inner and Outer Kinetochore. Curr. Biol. 21, 391–398. https://doi.org/10.1016/j.cub.2010.12.039

Shivaraju, M., Unruh, J.R., Slaughter, B.D., Mattingly, M., Berman, J., Gerton, J.L., 2012. Cell-Cycle-Coupled Structural Oscillation of Centromeric Nucleosomes in Yeast. Cell 150, 304–316. https://doi.org/10.1016/j.cell.2012.05.034

Sieben, C., Banterle, N., Douglass, K.M., Gönczy, P., Manley, S., 2018. Multicolor single-particle reconstruction of protein complexes. Nat. Methods 15, 777–780. https://doi.org/10.1038/s41592-018-0140-x

Sochacki, K.A., Dickey, A.M., Strub, M.-P., Taraska, J.W., 2017. Endocytic proteins are partitioned at the edge of the clathrin lattice in mammalian cells. Nat. Cell Biol. 19, 352–361. https://doi.org/10/f9vxd5

Sun, X., Zhang, A., Baker, B., Sun, L., Howard, A., Buswell, J., Maurel, D., Masharina, A., Johnsson, K., Noren, C.J., Xu, M.-Q., Corrêa, I.R., 2011. Development of SNAP-tag fluorogenic probes for wash-free fluorescence imaging. Chembiochem Eur. J. Chem. Biol. 12, 2217–2226.

Szymborska, A., Marco, A. de, Daigle, N., Cordes, V.C., Briggs, J.A.G., Ellenberg, J., 2013. Nuclear Pore Scaffold Structure Analyzed by Super-Resolution Microscopy and Particle Averaging. Science 341, 655–658. https://doi.org/10.1126/science.1240672

Thevathasan, J.V., Kahnwald, M., Cieśliński, K., Hoess, P., Peneti, S.K., Reitberger, M., Heid, D., Kasuba, K.C., Hoerner, S.J., Li, Y., Wu, Y.-L., Mund, M., Matti, U., Pereira, P.M., Henriques, R., Nijmeijer, B., Kueblbeck, M., Sabinina, V.J., Ellenberg, J., Ries, J., 2019. Nuclear pores as versatile reference standards for quantitative superresolution microscopy. Nat. Methods 16, 1045–1053. https://doi.org/10/gf85h3

Ustinov, N.B., Korshunova, A.V., Gudimchuk, N.B., 2020. Protein Complex NDC80: Properties, Functions, and Possible Role in Pathophysiology of Cell Division. Biochem. Mosc. 85, 448–462. https://doi.org/10.1134/S0006297920040057

Valverde, R., Ingram, J., Harrison, S.C., 2016. Conserved Tetramer Junction in the Kinetochore Ndc80 Complex. Cell Rep. 17, 1915–1922. https://doi.org/10.1016/j.celrep.2016.10.065

Virant, D., Vojnovic, I., Winkelmeier, J., Endesfelder, M., Turkowyd, B., Lando, D., Endesfelder, U., 2021. Unraveling the kinetochore nanostructure in *Schizosaccharomyces pombe* using multi-color single-molecule localization microscopy. bioRxiv 2021.12.01.469981. https://doi.org/10.1101/2021.12.01.469981

Walstein, K., Petrovic, A., Pan, D., Hagemeier, B., Vogt, D., Vetter, I.R., Musacchio, A., 2021. Assembly principles and stoichiometry of a complete human kinetochore module. Sci. Adv. 7, eabg1037. https://doi.org/10.1126/sciadv.abg1037

Watanabe, R., Hara, M., Okumura, E., Hervé, S., Fachinetti, D., Ariyoshi, M., Fukagawa, T., 2019. CDK1-mediated CENP-C phosphorylation modulates CENP-A binding and mitotic kinetochore localization. J. Cell Biol. 218, 4042–4062. https://doi.org/10.1083/jcb.201907006

Wei, R.R., Al-Bassam, J., Harrison, S.C., 2007. The Ndc80/HEC1 complex is a contact point for kinetochore-microtubule attachment. Nat. Struct. Mol. Biol. 14, 54–59. https://doi.org/10.1038/nsmb1186

Wei, R.R., Sorger, P.K., Harrison, S.C., 2005. Molecular organization of the Ndc80 complex, an essential kinetochore component. Proc. Natl. Acad. Sci. 102, 5363–5367. https://doi.org/10.1073/pnas.0501168102

Winey, M., Mamay, C.L., O’Toole, E.T., Mastronarde, D.N., Giddings, T.H., Jr, McDonald, K.L., McIntosh, J.R., 1995. Three-dimensional ultrastructural analysis of the Saccharomyces cerevisiae mitotic spindle. J. Cell Biol. 129, 1601–1615. https://doi.org/10.1083/jcb.129.6.1601

Wisniewski, J., Hajj, B., Chen, J., Mizuguchi, G., Xiao, H., Wei, D., Dahan, M., Wu, C., Kadonaga, J.T., 2014. Imaging the fate of histone Cse4 reveals de novo replacement in S phase and subsequent stable residence at centromeres. eLife 3, e02203.

Yan, K., Yang, J., Zhang, Z., McLaughlin, S.H., Chang, L., Fasci, D., Ehrenhofer-Murray, A.E., Heck, A.J.R., Barford, D., 2019. Structure of the inner kinetochore CCAN complex assembled onto a centromeric nucleosome. Nature 574, 278–282. https://doi.org/10.1038/s41586-019-1609-1

Yan, K., Zhang, Z., Yang, J., McLaughlin, S.H., Barford, D., 2018. Architecture of the CBF3–centromere complex of the budding yeast kinetochore. Nat. Struct. Mol. Biol. 25, 1103–1110. https://doi.org/10.1038/s41594-018-0154-1

Yatskevich, S., Muir, K.W., Bellini, D., Zhang, Z., Yang, J., Tischer, T., Predin, M., Dendooven, T., McLaughlin, S.H., Barford, D., 2022. Structure of the human inner kinetochore bound to a centromeric CENP-A nucleosome. Science 376, 844–852. https://doi.org/10.1126/science.abn3810

Zhang, W., Lukoyanova, N., Miah, S., Lucas, J., Vaughan, C.K., 2018. Insights into Centromere DNA Bending Revealed by the Cryo-EM Structure of the Core Centromere Binding Factor 3 with Ndc10. Cell Rep. 24, 744–754. https://doi.org/10.1016/j.celrep.2018.06.068

Zinkowski, R.P., Meyne, J., Brinkley, B.R., 1991. The centromere-kinetochore complex: a repeat subunit model. J. Cell Biol. 113, 1091–1110. https://doi.org/10.1083/jcb.113.5.1091

